# DIETS: a simple and sensitive assay to measure and control the intake of complex solid foods, like high-fat diets, in *Drosophila*

**DOI:** 10.1101/2023.06.07.543033

**Authors:** Manikrao R. Thakare, Prerana Choudhary, Bhavna Pydah, Suhas Sunke, R Sai Prathap Yadav, Pavan Agrawal, Gaurav Das

## Abstract

The fruit fly *Drosophila melanogaste*r offers a powerful model to study how diet affects the body and brain. However, existing methods for measuring their food intake often rely on dyes or tags mixed with food, which can be inaccurate due to how the flies absorb and eliminate them. Capillary-based assays like CAFE directly measure consumption, but only work with liquids and shorten fly lifespan. Additionally, capillary assays are incompatible with delivering viscous foods like high-fat diets. Even solidified high-fat diets tend to be sticky death traps for flies. Another longstanding challenge for fly researchers is that dietary restriction in flies involves diluting food, leading to compensatory feeding. To address these shortcomings, we have developed DIETS, a sensitive feeding assay that can be implemented even in low-resource settings. DIETS eliminates the need for labels and directly weighs the solid food consumed by small groups of flies over extended periods of hours to weeks. It allows us to deliver precise amounts of food to flies and implement accurate dietary restrictions. Importantly, DIETS is compatible with studying energy-dense high-fat diets. Using DIETS, we observed that, unlike a high-sugar diet, an isocaloric high-fat diet did not improve the flies’ ability to withstand starvation, even though they consumed more calories and had higher fat deposition.

## Introduction

The quality, quantity, and timing of food intake have a profound impact on the overall health, fecundity, cognition, and longevity of an organism. Food intake is influenced by internal states that reflect a lack of specific nutrients (Bjordal et al. 2014; Ribeiro and Dickson 2010; Toshima and Tanimura 2012; Prasad and Hens 2018; Steck et al. 2018; Musso, Junca, and Gordon 2021; Stafford et al. 2012; Malita et al. 2022; Morton, Meek, and Schwartz 2014), or by internal states that result from stress and pathogenesis (Crompton 1984; Yau and Potenza 2013). Food intake is also influenced by innate and learned sensory inputs, that signal the presence of nutrients (Shiu et al. 2022; Weaver et al. 2023; Das, Lin, and Waddell 2016; Oh et al. 2021; Fantino 1984). Humans and fruit flies share deep evolutionary conservation in physiology, metabolism, and nervous system function (Ugur, Chen, and Bellen 2016; Jennings 2011; Yamaguchi 2018; Diop and Bodmer 2012; Kornberg and Krasnow 2000). Precise genetic tools in flies allow the investigation of the neuro-physiological mechanisms that underlie appetite control and the myriad effects that follow the ingestion of specific foods (Lin, Senapati, and Tsao 2019; Guo, Pan, and Gong 2019; Yoshihara and Ito 2012; Owald, Lin, and Waddell 2015; Pool and Scott 2014).

For all such studies, the ability to measure, or control feeding precisely is critical. A deep understanding of the underlying biology remains incomplete without knowing how much food is consumed. Currently, there are multiple assays for estimating feeding in flies. However, they all suffer from problems that prevent their use across the board for different dietary compositions. Specifically, such shortcomings include relying on proxy measurements of feeding using dyes and other tags, using liquid-only diets, incompatibility with longitudinal studies, and the need to sacrifice flies for measurement.

As flies consume only ∼ 0.5 - 2.5 microlitres of food in a day (William W. Ja et al. 2007; Wu et al. 2020; Deshpande et al. 2014; Carvalho, Kapahi, and Benzer 2005), many fly-feeding assays rely on food tagged with dyes, radiolabeled molecules, or short oligonucleotides (Shell et al. 2018, 2021; Park, Tran, and Atkinson 2018; Wu et al. 2020; Geer, Olander, and Sharp 1970). The amount of ‘tag’ ingested, and eventually excreted acts as a proxy for consumption. Each tag must be carefully screened to determine whether their taste, absorption, incorporation, and excretion influence measurement, especially when physiology changes with age or genotype (Marx 2015; Deshpande et al. 2014; Shell et al. 2018, 2021). Quantification of an ingested tag requires sacrificing flies. Only the EX-Q (excreta quantification) assay avoids sacrificing flies by chasing out all ingested dye with an extended feeding period on unlabeled food. Food intake was then estimated by dye quantification from the excreta, leaving the flies alive for longitudinal experiments (Wu et al. 2020).

Ethologically, flies likely feed on liquid, solid, and semi-solid food sources. However multiple tagged and direct assays have been developed for flies solely for measuring liquid diet intake (Sellier, Reeb, and Marion-Poll 2011; Qi et al. 2015; Murphy et al. 2017; Yapici et al. 2016). Their designs are based on the CApillary FEeder Assay (CAFE), where liquid food consumed from a microcapillary is directly quantified by the drop in meniscus (William W. Ja et al. 2007). Recent reports show that feeding liquid diets over days from capillaries could nutrient-deprive flies and affect both lifespan and fecundity (Park, Tran, and Atkinson 2018; Lee et al. 2008; Vigne and Frelin 2010; Deshpande et al. 2014).

*Drosophila* is being widely and effectively used to gauge the effect of high-fat or high-sugar diets (HFD and HSD respectively) on the physiology, metabolism, nervous system function and behavior (Birse et al. 2010; Laura Palanker Musselman et al. 2011; L. P. Musselman, Fink, and Baranski 2019; Van Dam et al. 2021; Vaziri et al. 2020; May et al. 2019; Yang et al. 2023; Liao, Amcoff, and Nässel 2020). Capillary-based liquid assays are likely incompatible with viscous, multi-ingredient laboratory diets, especially liquid high-fat diets. Even when agarified, solid high-fat diets tend to become sticky and trap flies on them (Diop, Birse, and Bodmer 2017), especially with rising temperatures. Such problems could be the reason why very few fly studies with high-fat diets have attempted to measure actual food intake, especially over long periods (Shi et al. 2021; Eickelberg et al. 2022).

Here, we describe a simple, and versatile feeding assay that directly measures the actual amount of solid food consumed by groups of flies. Dubbed DIETS (<u>D</u>irect Intake <u>E</u>stimation and <u>T</u>racking of <u>S</u>olid food consumption), the assay is easily adaptable and inexpensive to establish even in low-resource settings. DIETS can measure feeding over both short and long periods, typically from 30 minutes to 24 h. Further, by periodically switching flies to fresh DIETS vials, longitudinal feeding measurement can be performed over weeks. By placing an additional food cup in a DIETS vial, we could also monitor short- and long-term feeding preferences in flies between two distinct foods. Crucially, the small surface area of the food make DIETS perfectly suited for complex high-fat diets. Flies do not stick to the food and do not die even at higher temperatures. We have also modified DIETS to a flat arena setup which can be combined with videography to determine the position of flies during an experiment and potentially record its contact with food.

DIETS addresses a significant drawback of fly dietary restriction (DR) assays. In flies, DR is mainly achieved by diluting food or nutrients (Hodge et al. 2022; Partridge, Piper, and Mair 2005; Carvalho, Kapahi, and Benzer 2005; Piper and Partridge 2007). However, food dilution leads to compensatory feeding and could complicate the interpretation of the results (Carvalho, Kapahi, and Benzer 2005; Vigne and Frelin 2010). With DIETS, we demonstrate that flies could be restricted to feeding an exact amount of complex foods by offering a predetermined amount of food in the cups. Hence, DIETS would be extremely useful for DR and Time Restricted Feeding (TRF) protocols with complex diets.

The mechanisms by which food intake may alter fitness are not all well understood. We employed DIETS to assess how two distinct calorie-dense, isocaloric food sources, one rich in sugar and the other rich in saturated fats, could affect food/calorie intake and contribute towards starvation resistance (SR). Surprisingly, we found that while flies ingested considerably higher calories on a high-fat diet than on a high-sugar diet, starvation resistance was only enhanced with the latter. The size of bodily fat deposits has been strongly correlated with starvation resistance. Our findings indicate that energy intake and body fat reserves do not necessarily correlate positively with the flies’ response to starvation.

## Results

### Measurement of feeding by directly weighing food consumed

The DIETS assay quantifies solid food consumption in flies. Consumption is directly measured by weighing the amount of food offered in 4 mm diameter ‘cups’ to a group of flies at the start of an experiment and finally weighing the amount of food left over at the end of the experimental period (Fig. 1A, Fig. S1A, and Movie S1). A 0.75 % agar bed at the bottom of the vial minimizes evaporation and provides a separate water source. A mean food weight loss from a control diet (CD; see Table ST1) due to evaporation was calculated from evaporation control vials without flies and was subtracted from concurrent intake measurements (Fig. 1A, Fig. S1B). We typically estimated consumption in sated CS-Q flies over 2 to 24 h periods, after which experiments were either terminated, or flies were transferred to fresh DIETS vials for further measurements (Fig. 1A). All experiments were carried out at 25 °C and 65 - 70 % relative humidity unless otherwise mentioned.

**Figure 1.**
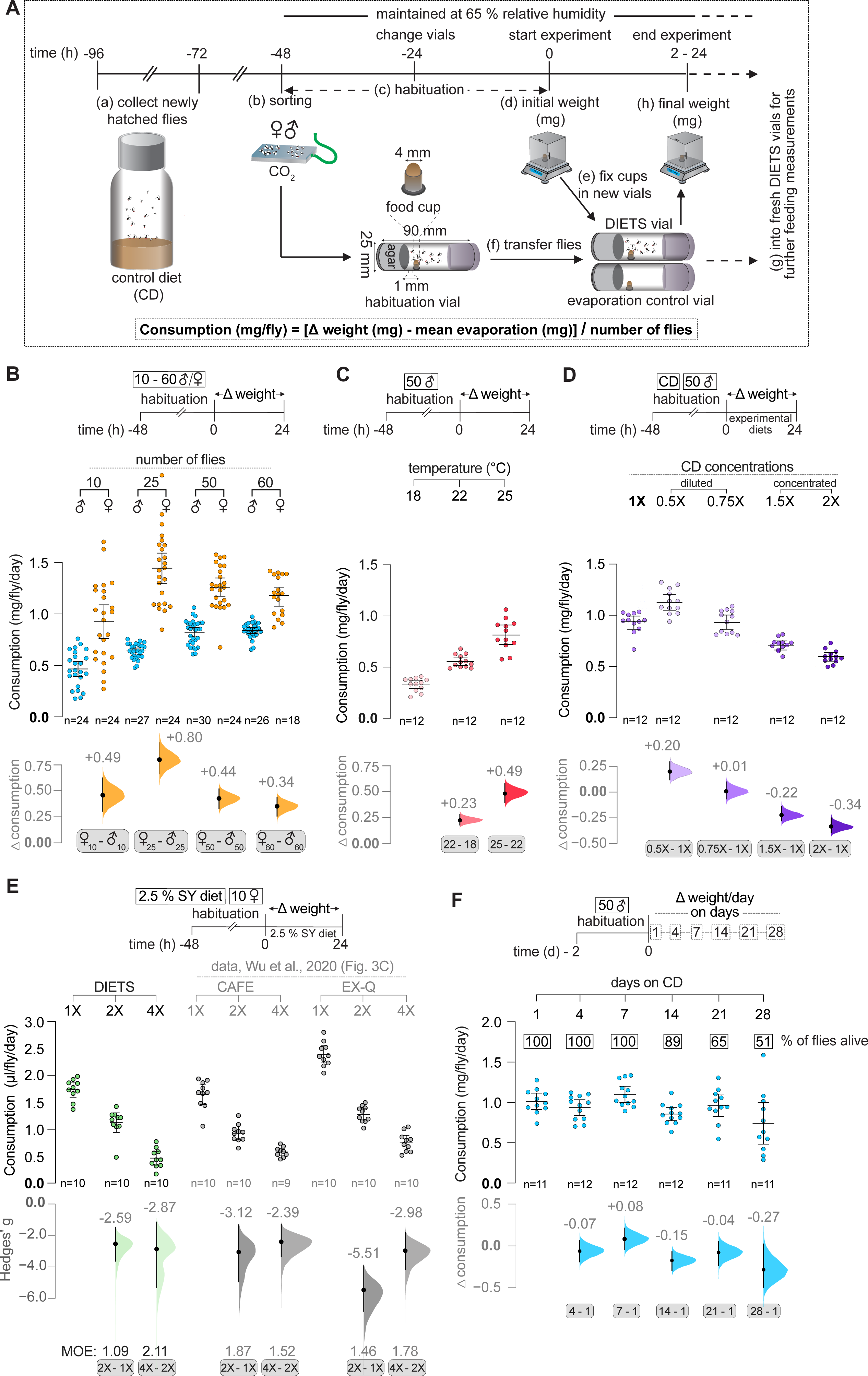
DIETS (<u>D</u>irect Intake <u>E</u>stimation and longitudinal <u>T</u>racking of <u>S</u>olid food consumption) is a sensitive assay that can measure food consumption in flies over days. **(A)** Schematic outline of the DIETS assay. **(B)** Comparison of 24 h control diet (CD) consumption in males and females of increasing group sizes. Females consistently eat a greater amount of food than males, regardless of the group size. Mean consumption by both sexes increased with group size. **(C)** CD consumption by groups of 50 male flies was measured over 24 h at different temperatures. An increase in the amount of food consumed was observed with an increase in temperature. **(D)** Feeding by groups of 50 male flies on varying concentrations of CD was measured. Flies show compensatory feeding upon diluting or concentrating CD. **(E)** Comparing the sensitivity of DIETS, CAFE, and EX-Q assays in resolving food intake difference between 1X, 2X, and 4X concentrations of 2.5 % sucrose - 2.5 % yeast extract (2.5 % SY) diet. DIETS show comparable resolution to CAFE and EX-Q. **(F)** Longitudinal measurement of CD consumption over days 1, 4, 7, 14, 21, and 28 for 50 male flies. Consumption mostly remains unaltered, except for a dip in feeding by the end of 28 days, possibly due to the mortality observed. Flies start dying between the 7^th^ and the 14^th^ day. Scatter plots of raw data with mean ± 95 % CI are shown. Δ ‘effect size’ plots below each raw data graph depict mean differences (black dots; values labeled) between relevant groups or standardized mean difference, hedge’s g for Fig. 1E. The 95 % CI (black lines) and distribution of the mean differences (curve), generated are also shown. See Materials and Methods for further details.

We empirically determined the size of the cups and the distance from the agar bed at which loss of food by evaporation from CD was minimal. Evaporation from a 4 mm diameter cup was lower than that from 6 mm or 8 mm diameter cups, and all were placed 1 mm away from the agar bed (Fig. S1C). Evaporation loss from a 4 mm diameter cup was minimal when placed 1 mm away from the agar bed and went up considerably when the distance of the cup from the agar bed was increased up to 5 mm at both 18 and 25 °C. As expected, evaporation loss was higher at 25 °C than at 18 °C (Fig. S1D).

Intuitively, food consumption should not change significantly with small changes in food cup position. We observed that the calculated CD consumption dropped if we increased the distance of food cups from the agar bed from 1 mm to 3 mm, or 5 mm when subtracting mean evaporation loss (Fig. S1E; left panel). However, the mean consumption values remained unchanged when the mean evaporation loss was not accounted for (Fig. S1E; right panel). This suggested that we were perhaps overestimating evaporation loss using the evaporation control vials, especially when cups were placed a little further away from the agar bed. As this overestimation of evaporation loss was the minimum at 1 mm (Fig. S1D), we decided to keep cups at this distance from the agar bed, unless otherwise required.

### DIETS is a sensitive and precise feeding assay

To characterize the resolving power and consistency of the DIETS assay, we measured feeding on CD after varying sex, group size, temperature, and food dilutions. Such conditions have been reported to have affected consumption (Wu et al. 2020; Klepsatel, Wildridge, and Gáliková 2019; William W. Ja et al. 2007; Deshpande et al. 2014). Firstly, males and females were segregated into groups of 10 to 60 flies to measure food consumption over 24 h at 25 °C. Like previous reports (Deshpande et al. 2014; Wu et al. 2020; Carvalho et al. 2006), the mean consumption of mated females was between 40 % to 125 % more than males from the same group size (Fig. 1B; see effect sizes or Δ consumption between two groups). We also observed that the consumption rate per fly increased with the group size and eventually plateaued with a group size of 50 for males and 25 for females (Fig. 1B, Table ST2). Therefore, as reported earlier, the alteration of social interactions with group size seemed to influence individual feeding behavior (Wong et al. 2009). As expected, the total food consumed by a group increased with the group size for both sexes and was clearly discernible with DIETS (Fig. S2A, S2B, and Table ST2). In experiments with mated females, we observed extensive egg deposition, mostly on the agar bed (Fig. S2C). Only ∼ 2 - 4 eggs were found on a food cup on average, with the number going up to ∼ 8 eggs for 60 females (Fig. S2D). As the mean egg weight was ∼ 32 □g (data not shown), egg weight could be ignored, especially if the ones on the outside of the cup were cleared before final weighing. In case larvae are seen on food, we suggest switching food more frequently and performing the experiments at a lower temperature if possible.

A recent study in flies has shown that an increase in temperature enhances metabolic rate and feeding (Klepsatel, Wildridge, and Gáliková 2019). We also saw a clear increase in CD consumption by male flies over 24 h when we raised the temperature from 18 °C to 22 °C and then from 22 °C to 25 °C (Fig. 1C; see effect sizes). Consumption on CD was even higher at 32 °C (Fig. S6C). We could detect such changes in temperature-dependent consumption with a sample size of 12 vials per condition.

Existing solid food feeding assays can detect compensatory increase or decrease in feeding in flies upon diluting or concentrating food (Deshpande et al. 2014; Wu et al. 2020). With DIETS, we were able to discern clear differences in feeding between 1X CD and 0.5X CD (∼ 21 % increase) or between 1X and 2X CD (∼ 56 % decrease) (Fig. 1D).

To directly compare the sensitivity of DIETS with the previously reported CAFE and EX-Q data (Wu et al. 2020), we measured compensatory feeding from 1X to 2X and to 4X concentrations of 2.5 % sucrose - 2.5 % yeast extract ( 2.5 % SY: 1X) diet (n=10, to match with Wu et al. 2020). We compared the standardized effect sizes in Hedge’s g along with its MOE (half the magnitude of 95 % CIs of the Hedge’s g: see Materials and Methods for further details) across assays (Fig. 1E). Between 1X and 2X diets, the analysis showed that DIETS and CAFE report effect sizes that are close to each other (−2.59 for DIETS, and −3.12 for CAFE). However, a larger effect size is seen with EX-Q (−5.51), mainly because 1X consumption measured by EX-Q was elevated (Fig. 1E). Between 2X and 4X diets, all three assays report quite similar effect sizes (−2.87 for DIETS, −2.39 for CAFE, and −2.98 with EX-Q). Keeping in mind the small size of the data sets (n=10), the MOE was also generally comparable (Fig. 1E). We concluded that DIETS was as sensitive as CAFE and EX-Q and likely equally precise (compare MOEs of all DIETS data reported in Table ST2).

To test whether DIETS can be used to assess longitudinal feeding, we transferred groups of 50 male flies into fresh assay vials every 24 h for 4 weeks. Actual consumption was assessed over the 1^st^, 4^th^, 7^th^, 14^th^, 21^st^, and 28^th^ day (schema; Fig. 1F). Feeding was reliably measured over the whole period, with very little change in feeding when compared to the 1st-day measurement, except on the 28^th^ day when a considerable feeding decline was observed. We also observed that with a large group size of 50, flies started to die around the 14^th^ day, and by the 28^th^ day, almost 50 % of the flies had perished (Fig. 1F).

Next, to test whether flies were starving in DIETS due to a smaller food source and hence eventually dying early, we compared the starvation resistance of male flies (10, 25, and 50 flies/group) after being in DIETS vials or normal food vials for 1 day and 7 days. We found that flies showed almost similar starvation resistance across group sizes and modes of food delivery (Fig. S3A). Together, our results suggest that no discernible change in survival is observed in DIETS up to 7 days. We also measured feeding on CD in two other strains available in the lab, *CS-BZ,* and *w^1118^*, and observed that *CS-BZ* had elevated feeding as compared to the other two strains (Fig. S3B).

To demonstrate the suitability of DIETS with thermogenetic manipulations at 32 °C, we activated Piezo-GAL4 labeled neurons using a temperature-sensitive cation channel, dTRPA1 (Hamada et al. 2008) during a 30-minute feeding period at 32 °C. Piezo-GAL4 labeled mechanosensory neurons densely innervate the crop and anterior midgut and regulate ingestion volume by controlling the crop size (S. Min et al. 2021). As reported earlier (S. Min et al. 2021), we too observed a defect in feeding, in fact, a more severe one, upon artificial activation of Piezo-GAL4 neurons compared to parental controls (Fig. S3C; left panel). Feeding reduction in Piezo-GAL4>UAS-dTRPA1 flies was not observed at 21 °C (Fig. S3C; right panel). Activation of serotonergic neurons labeled by R50H05-GAL4 induces hunger and feeding in sated flies (Albin et al. 2015). We replicated these results with DIETS, with increased feeding observed in R50H05-GAL4>UAS-dTRPA1 flies at 32 °C (Fig. S3C; left panel), but not at 21 °C (Fig. S3C; right panel).

### Short-term feeding measurements with DIETS

Next, we wanted to determine whether DIETS could be used for short-term feeding measurements. For this, we starved groups of 50 male flies on 0.75 % agar for 6 h, 12 h, or 24 h. We then measured the consumption of these differentially starved groups for 30 minutes in DIETS vials with Brilliant Blue FCF labeled CD (Fig. 2A, schema). Parallelly, flies were also processed after 30 minutes to measure consumption colorimetrically. Both techniques could detect the linear increase in consumption with increasing starvation and also detect sizable effect size differences with each step of starvation. For the sake of comparison across assay types, here too, we plotted standardized effect sizes in the form of Hegde’s g ( Fig. 2A). With DIETS, we had larger Hedge’s g values (Fig. 2A) than with blue dye incorporation across all comparisons (feeding after 6 -vs.- 0, 12 -vs.-6 and 24 vs. 12 h starvation times). The MOEs of these effect sizes are also comparable across the two assays (Fig. 2A; see values represented as effect sizes, MOEs; also in Table ST2). These results suggested that for short-term measurements, DIETS was more sensitive in picking differences than blue dye incorporation but equally precise.

**Figure 2.**
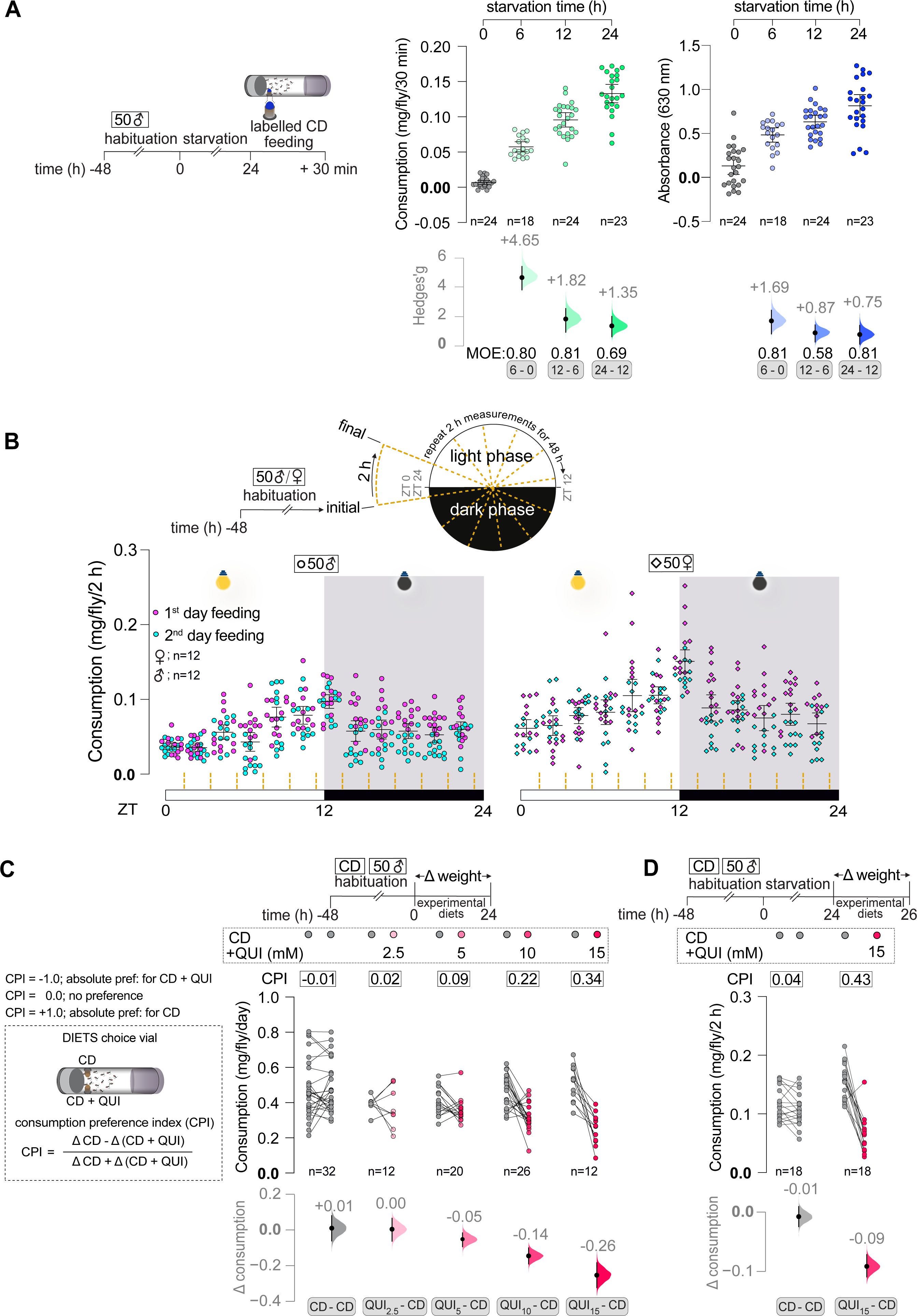
DIETS for short-term feeding measurements and food choice. **(A)** Comparing the sensitivity of DIETS with blue dye uptake in measuring short-term feeding. 30 minutes of feeding on CD supplemented with Brilliant blue FCF in differentially starved male flies was measured using DIETS. Flies showed an increase in feeding with increased starvation (left panel). A simultaneous dye accumulation measurement was conducted by crushing the flies after feeding (right panel). Hedge’s g is effect size, in this case, the difference in mean of two groups, standardized over the pooled standard deviation of the two groups. DIETS assay yields larger ‘g’ values compared to blue dye estimation for comparison of feeding between the same groups. **(B)** Continuous 2 h feeding measurements in sated male and female flies measured over 2 days. Both sexes show a distinct evening feeding peak just around the lights-off time at ZT = 11.5-12.5. **(C)** 24 h, food preferences between CD and CD mixed with quinine. Flies show an increasing dose-dependent aversion towards CD adulterated with quinine. **(D)** 2 h food preferences between CD and CD adulterated with 15 mM quinine in 24 h starved flies. Flies showed a strong aversion towards quinine-laced food. Scatter plots of raw data with mean ± 95 % CI are shown. Δ ‘effect size’ plots below each raw data graph depict mean differences (black dots; values labeled), between the two relevant groups being compared. The 95 % CI (black lines) and distribution of the mean differences (curve) generated by bootstrap resampling of the data are also shown. In (A) standardized effect size, Hedge’s g has been plotted, and MOEs have been reported. See Materials and Methods for further details.

Next, to demonstrate whether we can use DIETS for repeated short-term feeding measurements, we measured feeding every 2 h in groups of 50 male and female flies for 2 days continuously (Fig. 2B, schema). Previous studies had reported two peaks of feeding around the morning and evening light-dark transitions (May et al. 2019; Niu et al. 2021; Ro, Harvanek, and Pletcher 2014; Li et al. 2021; Xu, Zheng, and Sehgal 2008). However, we observed only a single evening feeding peak around ‘lights-off’ (Fig. 2B). Combining consumption measurements of the two ‘peak’ groups spanning the ‘lights off’ time and comparing them with similarly merged ‘day’ and ‘night’ groups on either side, we could see the effect of day to night transition on feeding (Fig. S4A). We also observed that otherwise, feeding levels were quite similar both during the day and night. From the above experiments, we concluded that DIETS is also sensitive and precise for both discrete and repeated short-term food intake measurements in sated flies.

### Short-term and long-term two-choice food preference with DIETS

We reasoned that dietary choice between two foods could also be measured if flies were given access to two cups of distinct foods. To validate food preference, we arranged choices between CD and CD laced with increasing concentrations (2.5 mM to 15 mM) of bitter-tasting quinine (QUI), for groups of 50 males, for 24 h (Fig. 2C, schema). Cups were placed 1 mm from the agar bed and diametrically opposite to each other. Fully assembled DIETS choice vials were covered with white paper to act as a diffuser and prevent light bias (Fig. S4B; images). In control DIETS vials, flies ate equally on average from both the cups with CD. As expected, flies preferentially fed from the CD cup over the CD + QUI cup, with increasing concentrations of quinine (Fig. 2C). However, combined consumption of food from both cups remained unchanged across quinine concentrations (Fig. S4C). Similarly, we measured short-term preference over 2 h in starved flies. We observed a strong preference to feed from the CD cup over the CD + 15 mM QUI food cup (Fig. 2D). Total consumption stayed constant in this case, as well (Fig. S4D). Food preference in DIETS can also be represented as a consumption preference index (CPI) (Fig. 2C; see inset). Corresponding CPI values are shown on top of the consumption plots (Fig. 2C and 2D).

### Implementing precise DR and TRF regimens with DIETS

Dietary restriction (DR) in flies is usually carried out by diluting food or protein components (Partridge, Piper, and Mair 2005; Bass et al. 2007; Piper and Partridge 2007; W. W. Ja et al. 2009; K.-J. Min and Tatar 2006; Hodge et al. 2022; Katewa et al. 2012). Since we could control the amount of food we offer to flies, we decided to ascertain whether we could use DIETS to achieve precise DR.

In the ‘no DR’ control groups, 56 mg of CD was offered to groups of male flies on average (Fig. 3A). We had determined that this amount of food was non-limiting to a group of 50 male flies (see Material and Methods for details). To implement 90 % DR, the mean food offered to a group of flies was adjusted to be ∼90-91% of 56 mg (51 mg), on average to groups of males (Fig. 3A; labeled ‘a’). We similarly aimed for various degrees of DR up to 60% [Fig. 3A; (a / 56) * 100. Ultimately, achieved DR was calculated based on average actual consumption in each group (Fig. 3A; labeled ‘b’), in proportion to the mean consumption (43 mg) in the ‘no DR’ group [Fig. 3A; (b / 43) * 100]. We found that the amount fed by groups of flies at all the DR levels clustered narrowly. This was also reflected in the extremely narrow MOEs of each group (Fig. 3A, see error distribution of effect sizes. Also see Table ST2). We concluded that the DIETS setup can be used to restrict the amount of food consumed by a group of flies within a few percentage points.

**Figure 3.**
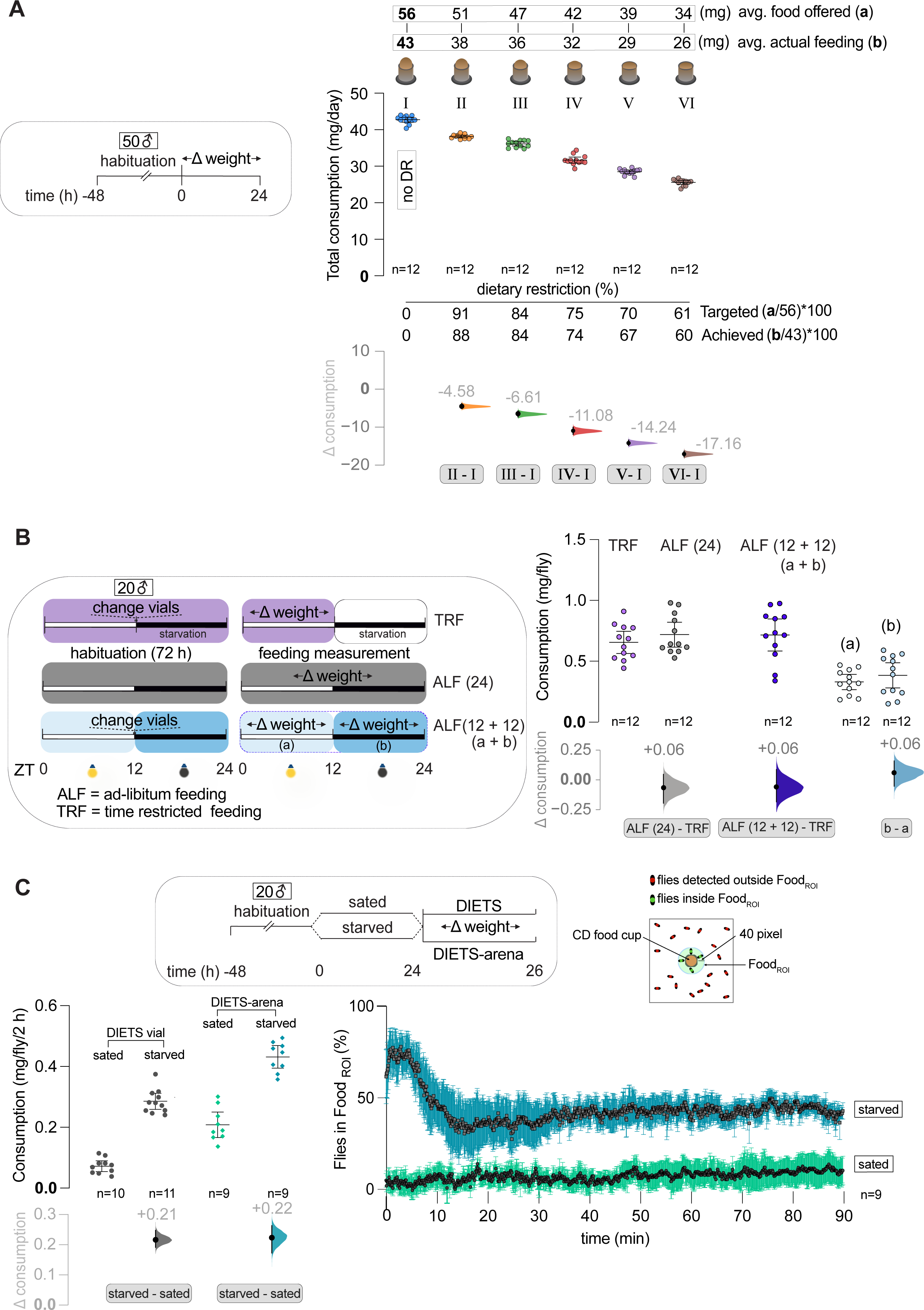
DIETS is well suited to study longitudinal feeding on high-fat diets and associated transcriptional and physiological changes. **(A)** In a longitudinal feeding study, flies were fed 5%, 10% and 20% saturated high-fat diets made with coconut oil (sHFD), and feeding measurements were conducted on days 1, 4, and 7. While consumption remained mostly constant up to 10 % extra coconut oil in the diet, feeding decreased noticeably with the addition of 20 % coconut oil. **(B)** Differential expression of *takeout* (*to*)*, Cytochrome P450-4e3* (*Cyp4e3*), *Odorant-binding protein 83a* (*Obp83a*), and *Turandot A* (*TotA*) was examined by qPCR from heads of flies that have been fed with 20 % sHFD for 1, 4, and 7 days, with respect to control flies fed CD. These genes were earlier found to be some of the most differentially expressed genes from an earlier study with a similar design. The direction and magnitude of differential expression of the above candidate genes on day 4 and 7 were comparable to that in day 7 samples from the previous study (Stobdan et al. 2019), as shown. **(C)** Energy intake was determined over 24 h periods on days 1, 4, and 7 in groups of flies fed 30 % HSD or isocaloric 10 % sHFD. The corresponding mean feeding values are represented above inside pink boxes. Flies fed on 10 % sHFD showed increased energy intake compared to the CD and HSD fed groups on all days. **(D)** The same flies were transferred to 0.75 % agar vials after 7 days of dietary treatment and the number of dead flies recorded every 6 - 12 hours. Survival time (left panel), and percent survival under starvation (right panel) of flies are shown. While HSD treatment increases starvation resistance, sHFD treatment does not. Scatter plots of raw data with mean ± 95 % CI are shown. For Figure 4C starvation resistance was plotted as Kaplan-Meier graph. Δ ‘effect size’ plots below each raw data graph depict mean differences (black dots; values labeled), between the two relevant groups being compared. The 95 % CI

Time-restricted feeding (TRF) protocols have been imposed on flies to gauge the effect on the overall health and longevity of flies (Cabrera, Young, and Axelrod 2020; Gill et al. 2015; Ulgherait et al. 2021). To investigate the impact of a daily pattern of feeding and fasting on food intake with DIETS, we measured CD consumption in groups of 20 male flies for 24 h under three different conditions: (a) the TRF group where feeding was restricted to ‘lights-on’ phase, ZT = 0 - 12, (b) An *ad libitum* feeding “ALF (24)” group with food cup changed every 24 h at ZT = 0, and (c) A second a*d libitum* feeding group, “ALF (12 + 12)” with food cup changed every 12 h at ZT = 0 and 12. All groups were habituated as described above for 3 days to let them adjust to their respective conditions. Consumption was measured for the next 24 h period (Fig. 3B; schema). In agreement with previous reports (Cabrera, Young, and Axelrod 2020; Gill et al. 2015), we found that flies under TRF consumed similar amount of food during the daytime as the ALF (24) and ALF (12 + 12) groups managed to eat over a whole day (Fig. 3B). Next, upon comparing daytime feeding ‘a’ with nighttime feeding ‘b’ within the ALF (12+12) group, our data revealed that flies fed equally during the day and night (Fig. 3B). We concluded that DIETS can be used to implement TRF protocols with solid foods and simultaneously measure feeding too.

### Combining consumption measurement with video tracking: DIETS-arena

DIETS vials are not ideal for videographic analysis because of their dimensions and also due to the location of the cups next to the agar bed. As a proof of concept that video analysis of fly location can be combined with DIETS style measurement of food consumption, DIETS-arenas were designed (Fig. S5A). First, to determine the efficacy of the flat DIETS-arenas, we measured 2 h CD consumption in sated and starved male flies with both DIETS vials and DIETS-arena side by side (Fig. 3C; schema). We observed a similar increase in feeding in the starved groups compared to the sated group with both vials and arenas (Fig. 3C, left panel). The feeding baseline was higher in the DIETS-arenas compared to DIETS vials for both sated and starved flies. Further investigation will be needed to determine the cause. Next, to analyze the temporal dynamics of feeding with the same set of sated and starved flies, we defined a Food_ROI_ around the food cup in the center of the arena (see Material and Methods). We found that the percentage of starved flies that occupied the Food_ROI_, went from 74 % to ∼ 40 % in the first 20 minutes. Thereafter starved flies maintained an average of ∼ 44 % Food_ROI_ occupancy. In contrast, fed flies maintain a low and steady ∼ 11 % occupancy of the Food_ROI_ over 90 minutes (Fig. 3C, right panel). This pattern is also reflected in temporal snapshots from two representative sets, each of sated and starved flies (Fig. S5B). In conclusion, with the help of video analysis from DIETS-arenas, we could visualize dynamic changes in food occupancy that could be correlated with the increased feeding observed in starved flies.

### Longitudinal measurements of high-fat diet (HFD) intake with DIETS

High-fat diets (HFDs) for flies have been typically made by adding coconut oil as a source of saturated fat to fly food (Birse et al. 2010; Diop, Birse, and Bodmer 2017). HFDs are difficult to work with as the fat melts even at ∼ 25 °C, rendering the food sticky and lethal for flies unless care is taken (Diop, Birse, and Bodmer 2017). The sticky consistency of fatty food could potentially affect feeding too. While a previous study claims the use of CAFE for delivering high-fat food (Shi et al. 2021), we sought to independently verify the suitability of CAFE with 10 % sHFD made in the lab. We re-confirmed earlier observations in the lab that such complex food tends to separate within the CAFE capillaries, leading to gaps in the continuous supply of food (Fig. S6A). We reasoned that DIETS, with its low food surface area, may resolve such issues. We prepared HFDs with added 5 % to 20 % coconut oil at the expense of water in the CD. We recalibrated evaporation control vials for HFDs and found that HFD food cups were best placed 2 mm from the agar bed. At lesser distances (1 mm), the HFD tended to accumulate moisture, leading to misleading changes in weight (data not shown). Consumption was measured on the 1^st^, 4^th,^ and 7^th^ day (Fig. 4A, schema). We found that consumption of diets with 5 % and 10 % added coconut oil was comparable with CD throughout (Fig. 4A). As the added lipids made the food more energy-rich, energy intake was increased at both these conditions compared to CD (Fig. S6B). We also saw that consumption of the diet with 20 % added oil was considerably reduced from all the other groups (Fig. 4A; see effect sizes HFD). As a result, energy intake in the 20 % HFD group was virtually similar or slightly increased to the 10 % HFD group (Fig. S6B). To determine whether decreased food intake on HFDs reflects an energy limit sensing mechanism or simply because higher fat concentrations are taste aversive to flies requires further investigation. DIETS was also suitable for consumption measurement on HFD even at 32 °C (Fig. S6C), enabling thermogenetic experiments with HFDs. Overall, we concluded that DIETS was suitable for feeding measurements on HFDs over a wide range of temperatures.

**Figure 4.**
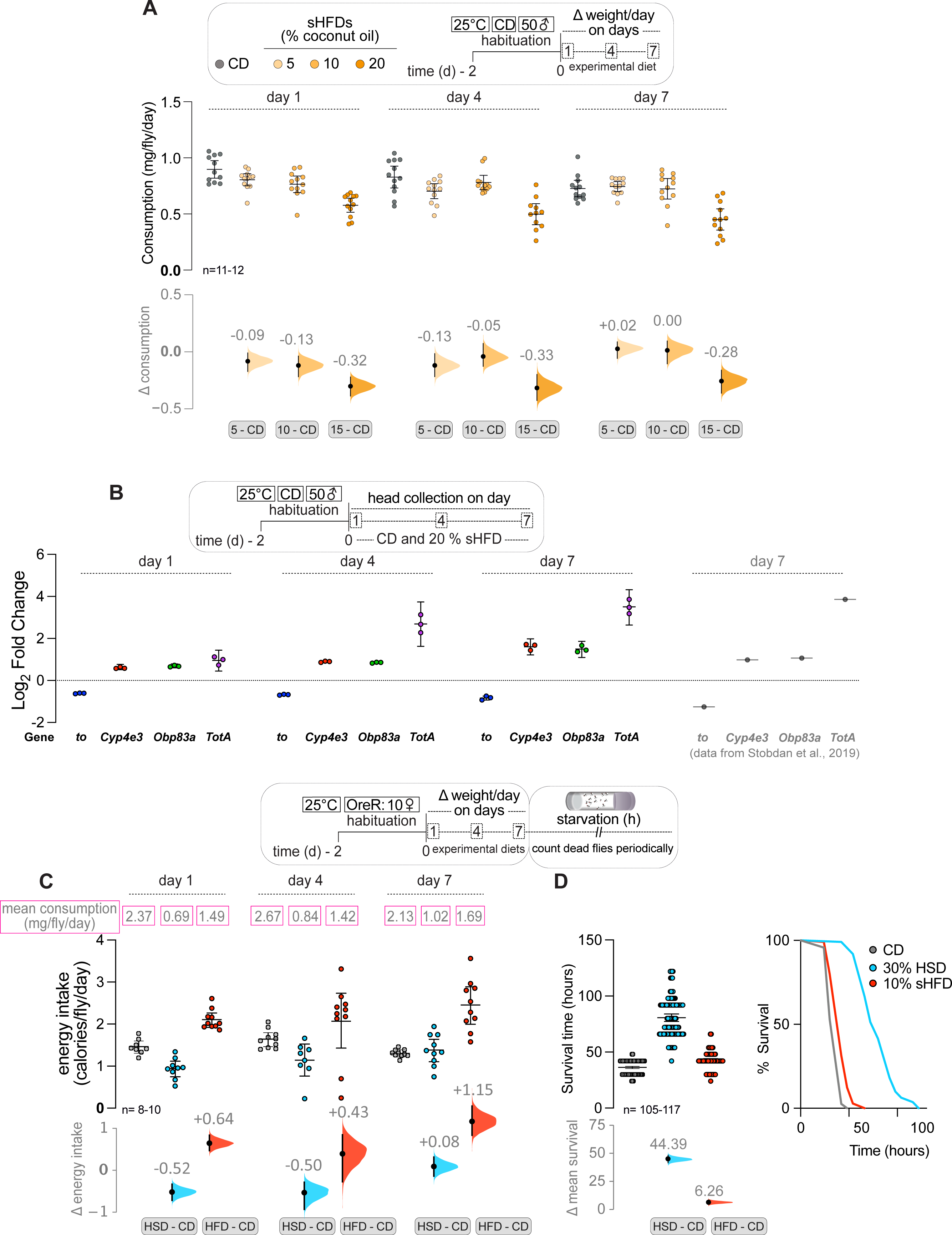
DIETS is well suited to study longitudinal feeding on high-fat diets and associated transcriptional and physiological changes. **(A)** In a longitudinal feeding study, flies were fed 5%, 10% and 20% saturated high-fat diets made with coconut oil (sHFD), and feeding measurements were conducted on days 1, 4, and 7. While consumption remained mostly constant up to 10 % extra coconut oil in the diet, feeding decreased noticeably with the addition of 20 % coconut oil. **(B)** Differential expression of *takeout* (*to*)*, Cytochrome P450-4e3* (*Cyp4e3*), *Odorant-binding protein 83a* (*Obp83a*), and *Turandot A* (*TotA*) was examined by qPCR from heads of flies that have been fed with 20 % sHFD for 1, 4, and 7 days, with respect to control flies fed CD. These genes were earlier found to be some of the most differentially expressed genes from an earlier study with a similar design. The direction and magnitude of differential expression of the above candidate genes on day 4 and 7 were comparable to that in day 7 samples from the previous study (Stobdan et al. 2019), as shown. **(C)** Energy intake was determined over 24 h periods on days 1, 4, and 7 in groups of flies fed 30 % HSD or isocaloric 10 % sHFD. The corresponding mean feeding values are represented above inside pink boxes. Flies fed on 10 % sHFD showed increased energy intake compared to the CD and HSD fed groups on all days. **(D)** The same flies were transferred to 0.75 % agar vials after 7 days of dietary treatment and the number of dead flies recorded every 6 - 12 hours. Survival time (left panel), and percent survival under starvation (right panel) of flies are shown. While HSD treatment increases starvation resistance, sHFD treatment does not. Scatter plots of raw data with mean ± 95 % CI are shown. For Figure 4C starvation resistance was plotted as Kaplan-Meier graph. Δ ‘effect size’ plots below each raw data graph depict mean differences (black dots; values labeled), between the two relevant groups being compared. The 95 % CI (black lines) and distribution of the mean differences (curve) generated by bootstrap resampling of the data are also shown. See Materials and Methods for further details.

As reported earlier (Birse et al. 2010), upon lipophilic Oil-Red-O staining, we detected increasing mid-gut fat deposition after 7 days with an increasing percentage of oil in the flies’ diet (Fig. S6D). This showed that HFD feeding in DIETS yielded expected physiological changes. Next, to ascertain whether feeding from HFDs in DIETS influenced gene expression changes in key genes, as reported earlier by RNA-Seq (Stobdan et al. 2019), we performed q-PCR analysis. We extracted RNA from the heads of male flies kept on either 20 % HFD or CD for 1, 4, and 7 days and probed gene expression using q-PCR (see Materials and Methods). We re-analyzed the raw RNA-Seq data generated previously (Stobdan et al. 2019) using DESeq2 (Love, Huber, and Anders 2014) and used p-adjusted < 0.05 as our threshold (Table ST3). From this analysis, we selected four of the most differentially expressed genes (DEGs) to analyze from our collected head RNA samples. Out of these, three were upregulated, namely *takeout* (*to*), *Obp83a*, and *Cyp4e3*, and one down-regulated gene, *TotA* (Fig. 4B). Our experiment revealed dynamic changes in the expression of the selected genes due to HFD feeding. On day 1, the expression of these genes showed variability as compared to the earlier study (Stobdan et al. 2019). However, by day 4 and then on day 7, their expression was consistent with Stobdan et al., 2019 (Fig. 4B). We concluded that DIETS enables us to measure feeding in a group of flies and subsequently use the same animals for post-feeding analyses like gene expression changes.

### A saturated high-fat diet fails to enhance starvation resistance

Chronic consumption of diets rich in sugar improves starvation resistance (SR) in female flies, likely due to increased bodily fat deposits (van Dam et al. 2020; Chandegra et al. 2017; Lee and Jang 2014; Strilbytska et al. 2022; Baldal, Brakefield, and Zwaan 2006; Ballard, Melvin, and Simpson 2008). However, although sHFDs increase body fat deposits (Fig. S6D), a solitary study has studied and reported a mild increase in SR upon sHFD feeding (Heinrichsen and Haddad 2012). As DIETS improve how sHFD is delivered to flies and likely decreases aversion to food due to sticky surface texture, we wanted to re-examine the role of sHFDs on SR. Hence, we provided OreR flies with an isocaloric 30 % high sugar diet (HSD: ∼21 % sucrose + other sugars) and 10 % sHFD (coconut oil) for 7 days. Feeding was assessed on days 1, 4, and 7. Subsequently, flies were subjected to starvation stress in 0.75 % agar vials and survival was ascertained every 24 h (schema; Fig. 4C). We found that flies on 30 % HSD, consumed significantly less food, but maintained a total energy intake comparable to flies on CD control food (Fig. S6E and 4C). In contrast, flies on 10 % sHFD, consumed amounts intermediated between those of flies on CD and 30 % HSD (Fig. S6E). However, sHFD being energy dense, the flies consumed significantly more energy than the control flies (Fig. 4C). Subsequently, we found that a 30 % HSD diet promotes increased SR in female flies as reported earlier. However, 10 % sHFD-fed flies showed near identical SR compared to control flies (Fig. 4D). We concluded that a 10% sHFD diet is unable to enhance SR in flies.

## Discussion

DIETS is a direct measurement of food consumed by flies with no added dyes or tracers (Fig. 1). Therefore, physiological changes associated with genotype or age, including changes in the gut, do not influence measurement. While DIETS is compatible with group sizes varying from ∼ 10 - 60 flies, we recommend that the group size should be optimized for specific experiments. When we measured food consumption in groups of 50 males, there was significant death after 2 weeks of feeding in DIETS vials (Fig. 1E). This could be due to increased stress and struggle for a small food source in larger groups, and likely to be mitigated by using a smaller group size for longitudinal measurements. Indeed, previous longevity studies have used much smaller group sizes of 10 - 15 flies (Lee et al. 2008; Metaxakis and Partridge 2013; Zanco et al. 2021; Krittika and Yadav 2020). We have shown that DIETS could be also utilized for measuring food preferences (Fig. 2). With DIETS we can easily monitor potential shifts in food preference between two, perhaps even three food sources, over days and weeks. The assay is also precise, and with proper humidity control, yields measurements with low variability (see MOEs in Table. ST2).

With regards to versatility, ease, and cost of use, only CAFE and EX-Q assays are comparable with DIETS (Fig. 1E). We show here that DIETS is as sensitive as CAFE (William W. Ja et al. 2007; Diegelmann et al. 2017) and EX-Q (Wu et al. 2020) to discern biologically relevant differences in food consumed (Fig. 1E; see effect sizes).

While CAFE directly measures liquid food intake, it has major drawbacks. Feeding long-term in CAFE significantly reduces both egg-laying and lifespan (Wong et al. 2009; Deshpande et al. 2014). Additionally, while studies have reported using CAFE for HFDs (Shi et al. 2021), we show that complex diets like HFDs separate inside capillaries, making it impossible to measure consumption (Fig. S6A). EX-Q is well suited for longitudinal studies, for use with complex diets, for estimating food preference with two compatible dyes, and possibly also for dietary restriction protocols (Wu et al. 2020; Shell et al. 2021). However, any protocol requiring measurement for short consecutive periods will not be possible with EX-Q because of the ∼ 3 h ‘chase’ phase on non-dyed food required to excrete out all the dye in flies. Therefore, longitudinal measurements with EX-Q require an extra fly transfer step and estimate daily consumption for only a 21 h period. Flies can independently regulate food and water intake (Van Dam et al. 2021; Fanson, Yap, and Taylor 2012; W. W. Ja et al. 2009). In DIETS vials, the agar bed provides a separate water source to allow flies to meet their water needs independently from their food drive (Fig. 1A). No separate water source is provided in the EX-Q assay (Wu et al. 2020). Ultimately, EX-Q is still an indirect method of feeding quantification, and any change in dye absorption or excretion with age or genotype would affect the measurement.

The DIETS assay is versatile and measures feeding over different time scales for diverse foods, including high-fat diets that are difficult to work with (Diop, Birse, and Bodmer 2017). This ability of DIETS is an especially significant feature as it allows long-term experiments with HFDs with accurate fatty food intake measurement. We are not aware of such studies in the current literature. One significant shortcoming of DIETS is that it cannot monitor feeding individuality in single flies. Capacitance-based single fly feeding assays like FLIC, flyPAD, optoPAD, or dFRAME are highly sensitive and outperform DIETS in determining feeding microstructure, including feeding duration and frequency of individual feeding bouts (Ro, Harvanek, and Pletcher 2014; Itskov et al. 2014; Moreira et al. 2019; Niu et al. 2021; Weaver et al. 2023). However, these assays do not measure feeding directly and only quantify the duration and frequency of fly proboscis contact with food. The contact of the proboscis with food may not always correlate directly with the amount ingested and may vary with age or genotype. Ultimately, these assays have much higher setup and repair costs than DIETS and are not affordable for many. Here, we demonstrate that direct food intake measurements can be carried out from flat and transparent DIETS-arenas and combined with video-based temporal analysis of a fly’s position. Similar to previous studies (Niu et al. 2021; Jaime et al. 2023; Weaver et al. 2023), with the right camera, lighting, and number of flies, the feeding microstructure of a single ‘tagged’ fly in a group can also be recorded from DIETS-arenas.

The dilution of food or its protein components leads to compensatory feeding in flies (Partridge, Piper, and Mair 2005; Chapman and Partridge 1996; Bass et al. 2007; Mair et al. 2003). To the best of our knowledge, DIETS is the only assay that can implement precise DR in flies without worrying about compensatory consumption (Fig. 4A). Additionally, like CAFE, time-restricted feeding (TRF) protocols (Cabrera, Young, and Axelrod 2020; Gill et al. 2015) are easily carried out with DIETS, with the added advantage of being able to work with high-fat and high-sugar foods. This ability to control and measure food intake will also make it possible to calibrate and deliver precise amounts of drugs or infectious agents to a group of flies.

Multiple lines of evidence indicate that increased fat deposits strongly correlate with increased SR in flies. Flies selected over generations for increased SR show enhanced body fat deposits (Schwasinger-Schmidt, Kachman, and Harshman 2012; Hardy et al. 2017; Chippindale, Chu, and Rose 1996; Harshman, Hoffmann, and Clark 1999). Foods with higher carbohydrate content (or lower P: C ratio) show increased bodily lipid content and increased SR (Lee and Jang 2014). As mentioned above, other studies on HSD and subsequent SR support the same idea (van Dam et al. 2020; Chandegra et al. 2017; Lee and Jang 2014; Strilbytska et al. 2022). We tested SR in OreR flies after a week-long exposure to isocaloric HSD and sHFD. We found that while HSD treatment expectedly improved SR over control, sHFD did not (Fig. 4D). This was despite higher energy consumption on sHFD (Fig. 4C) and likely increased fat deposit, similar to our observation for CS flies (Fig. S6D). We therefore speculate that feeding on HSD versus sHFD, leads to the formation of bodily fat deposits that could differ in aspects like composition, lipid droplet structure, and the rate of utilization during starvation. Ultimately, one or more of these factors could alter SR, and further investigations are needed to understand the exact mechanisms.

Our DIETS platform will be ideal for similar studies, where precise delivery of complex diets and accurate measurements of their consumption will be necessary. We anticipate that the ability of DIETS to precisely estimate and control food intake, its versatility, and the low cost of deployment will lead to its wide adoption.

## Materials and Methods

### A. Fly lines and rearing

We used the following fly strains for all experiments: Wild type Canton-S (CS-Q), Oregon-R (OreR), CS-BZ, and *w^1118^*, Piezo-GAL4 (BDSC # 59266) (S. Min et al. 2021), R50H05-GAL4 (BDSC # 38764) (Albin et al. 2015), and UAS-dTRPA1 (Hamada et al. 2008). All flies were reared on modified Bloomington’s semi-defined food (Laura Palanker Musselman et al. 2011) at 25 □ and 65 % humidity, with a 12 h:12 h light: dark cycle inside incubators. This food is referred to as the control diet (CD: see Table. ST1 for ingredients). Temperature-sensitive UAS-dTRPA1 flies were raised at 18 °C and transferred to 32 °C 30 minutes before the start of experiments to induce transgene expression.

### B. Chemicals and other materials

Food ingredients, namely brewer’s yeast (Prime brand, instant dry yeast), table sugar (Madhur brand, refined sugar) and cold-pressed coconut oil (Nutiva), Brilliant blue FCF food grade ( Vidhi Manufacturers 2490) were procured from local vendors. Other components, like agar (Himedia: GRM666-500G), peptone (Himedia: GRM001-500G), yeast extract powder (Himedia: RM027-500G), sucrose (MP biomedicals: 194018), propionic acid (Sigma-Aldrich: P5561-1L), methyl-4-hydroxybenzoate (Himedia: GRM1899-500G), MgSO_4_ X 7H_2_O (Sigma-Aldrich: M1880), CaCl_2_ X 2H_2_O (Sigma-Aldrich: 223506), absolute ethanol (Merck: 64-17-5), Oil-red-O (Sigma-Aldrich: 00625-100G), quinine hydrochloride dihydrate (Sigma-Aldrich: Q1125-5G), Triton X-100 (MP biomedical: SKU:02194854-CF) and paraformaldehyde (Sigma-Aldrich: P6148-500G), 0.22 □m syringe filter (MERK: Z260479) were procured from authorized vendors. Light filters used for uniform red light are 182: light red, 158: deep orange, and 810: zoom diffusion 1 (Bright Light International). The following chemicals were used for RT-PCR; Trizol (Invitrogen: 15596026), Zymo Direct Zol kit (Zymo Research: R2051), Zymo RNA Clean and Concentrator kit (Zymo Research: R1013), High-Capacity cDNA Reverse Transcription kit (Thermo-Fisher Scientific: 4368814), KAPA SYBR FAST Universal (Sigma-Aldrich: KK4602), QuantStudio™ 6 flex Real-Time PCR System (Applied Biosystems: 4485691).

### C. Diet preparation

To prepare the Control Diet (CD), ingredients (Table ST1) were weighed and mixed in 60 mL reverse osmosis (RO) filtered water. The mixture was then cooked in a microwave oven, with 2-3 intermittent stirring for ∼ 2 minutes. The food mix was then allowed to cool to 60 □ before adding methylparaben, propionic acid, CaCl_2,_ and MgSO_4_. The volume was adjusted to 100 mL by adding RO water. This mixture was then thoroughly stirred with a whisk. To prepare diets with 0.25X to 2X concentrations of CD (Fig.1D), we separately weighed all the ingredients for each concentration. Agar concentration was maintained at 1 % for all the dilutions. To prepare CD spiked with quinine (Fig. 3C), the required amounts of quinine were added to the molten CD, below 60 □ and mixed thoroughly by shaking or vortexing. 2.5 % sucrose - 2.5 % yeast diet ( 1X SY diet), 5 % sucrose - 5 % yeast diet (2X SY diet), and 10 % Sucrose - 10 % yeast diet (4X SY diet) were prepared as previously described (Deshpande et al. 2014; Wu et al. 2020). To prepare saturated high-fat diets (5 %, 10 %, and 20 % HFDs, w/v), the required amount of molten coconut oil was added to cooked food and then mixed with a whisk. The added coconut oil replaced an equal volume of water in the recipe. The food mix was allowed to cool to 60 □ and methylparaben, propionic acid, CaCl_2,_ and MgSO_4_ were added. The food was mixed thoroughly again with a whisk, and the volume was adjusted to 100 mL by adding RO water. A 10 % HFD was isocaloric to 30 % high-sugar diet (HSD). 9% sucrose is present in the CD and to prepare 30 % w/v HSD, an extra ∼ 21 % sucrose was added to replace the water in the CD. High-fat diets (HFDs) and high-sugar diets (HSDs) exhibited hygroscopic properties, leading to the accumulation of moisture on the food surface. We empirically determined the distance at which the food cups had to be fixed from the agar bed to prevent this. Typically it was between ∼ 1-5 mm.

### D. Feeding quantification by DIETS assay

#### i) DIETS vial preparation

Standard fly vials (inner diameter: 23 mm, outer diameter: 25 mm, height: 90 mm) and 200 µL microcentrifuge tube caps as food cups were assembled as follows. 7 mL 0.75 % agar solution was added to each vial to form an agar bed. After the agar solidified, it was allowed to dry for 12 to 18 h at room temperature. If used immediately, the agar layer leaks water, in which flies get trapped. An agar bed reduces the weight loss from food cups due to evaporation and provides a separate water source for flies to drink water. Solid food was provided in 200 µL microcentrifuge tube (PCR-02-C; Axygen) caps (food cups) of 4 mm diameter. For most experiments, ∼ 70 µL of molten food from stored aliquots were pipetted into food cups (Supplementary Fig. 1A). Once the food solidified, we noted the initial weight of the cup + food. The initial weight of the food cups (cup + food) was kept between a narrow range for consistency, typically ∼ 125 - 135 mg for CD. Freshly prepared food cups were fixed on the inside of the agar vials, normally at a distance of 1 mm from the agar, with a small square of double-sided tape. Evaporation control vials were prepared identically.

#### ii) Food consumption and calorie intake measurement

Flies were collected over the first day after eclosion, transferred to new food bottles, and allowed to age for one more day. Lines that carried transgenic constructs were aged for ∼ 8 - 10 days after initial collection. On the 3^rd^ day (∼ 9^th^ - 11^th^ day for transgenic lines), flies were segregated by sex under CO_2_ anesthesia. Male or female groups of 50 flies were placed in prepared DIETS vials and habituated for two further days before actual consumption measurement. Flies were transferred to fresh DIETS vials once after 24 h during the habituation process (Fig. 1A).

Fly groups were then transferred to DIETS vials with pre-weighed food cups for feeding measurements. Evaporation control vials with no flies was maintained alongside (Fig. 1A and Supplementary Fig. 1A). Viscosity of a specific food influences the rate of evaporation. We also observed that certain viscous foods, like foods with high fat and high sugar, in cups placed at 1 mm from the agar bed tended to absorb moisture. Conversely, in diluted, less viscous foods, evaporation loss increased. Therefore, with each new diet, we empirically determined the position of the food cups from the agar bed to prevent moisture accumulation or loss by evaporation in evaporation control vials below 10 % of the total food consumed. The humidity level of the incubator could also be adjusted to keep evaporation below the desired level. Most experiments were carried out at 25 □ and 65 % humidity.

Flies were transferred to fresh DIETS vials at the end of the period of feeding measurements. The reduction in food cup weight (food consumed + evaporation loss) was determined. The mean value of evaporation-mediated loss in weight (from 4 - 6 control vials) was deducted from the reduction in food cup weight to derive the total food consumed by a group of flies. This value was usually plotted as an average consumption per fly over time (e.g., mg/fly/day) or sometimes as total consumption for a group of flies (mg/day).

Energy intake (in Fig. 4C and S6B) was calculated from the amount of food consumed on CD and HFD, using the calorie values given in (Laura Palanker Musselman et al. 2011) and represented as calorie/fly/day. The caloric value of the Nutiva coconut oil was taken from the product label.

#### iii) Feeding measurement in starved flies with DIETS and blue dye uptake

Food intake in starved flies was measured by DIETS assay, and intake was parallelly validated using a spectrophotometric estimation of Brilliant blue FCF dye. To measure feeding in starved flies, 2 - 3 day old male flies were habituated in DIETS vials for 2 days. For starvation, flies were kept in 0.75 % agar vials for specified times. Flies were then transferred to DIETS vials with pre-weighed 1 % Brilliant blue FCF 1 dye-labeled CD and allowed to feed for 30 minutes. Flies were homogenized in 500 µl of PBST (1x PBS + 1 % Triton X-100) using a hand-held homogenizer (BenchTop Lab Systems: BT-MT-13K). Homogenates were centrifuged at 4,000 g for 5 minutes, and 100 µl of cleared supernatant was loaded into a 96-well plate. Sample absorbance was measured at 630 nm on a multi-mode plate reader (SPECTRAmax 250, Molecular Devices). Absorbance from a control set of flies fed with food without dye was used to determine the baseline absorbance of fly lysates (blank).

#### iv) Round-the-clock circadian feeding measurements

2 - 3 day old male and female flies were habituated in DIETS vials for two days. Feeding measurement was initiated at 12 noon [Zeitgeber time (ZT) = 3.5; our light-on time, or ZT = 0, was 8:30 AM] in fresh vials. Flies were then transferred to fresh DIETS vials every 2 h for the next 50 h, and continuous measurements were taken. The first 2 h feeding measurement was discarded, as it showed spiked feeding, likely due to the fact that they had been transferred to fresh food after 24 h. Fly transfers during the 12 h dark phase were carried out in red light.

#### v) Two food choice feeding measurements with DIETS

To measure food preference over varying time scales, two cups with distinct foods were placed 1 mm from the agar bed, diametrically opposite each other, inside DIETS vials. During the experiment, the two cups were kept parallel to the tray, facing each other. To avoid a bias due to uneven lighting, individual vials were wrapped in an 8.5 * 8.5 cm white printing paper sheet, acting as a light diffuser (Fig. S3A). Before measuring feeding from two food sources, flies were habituated in DIETS choice set up, with two CD cups for 48 h. Flies were starved, if the experiment demanded, after the habituation step. A consumption preference index (CPI) was also calculated as CPI_Food A_ = [(feeding from Cup A - feeding from Cup B) / total consumption from both the food cups].

#### vi) DR protocol with DIETS

We have achieved DR in flies by offering a fixed amount of diets in food cups to a group of flies for 24 h, after 2 days of habituation on the CD diet. As a ‘NO DR’ control (*ad-libitum* feeding), an average of 56 mg of CD was offered to groups of 50 male flies. This value was derived from summing the mean consumption of 50 male flies (∼ 39 mg: Fig. S2A), mean evaporation from food cup (∼ 4 mg: Fig. S1C), and mean leftover food (∼ 10-13 mg: data not shown). To implement increasing degrees of DR (∼ 90 % to 60 %), we gradually reduced the mean food offered in cups by ∼ 3 to 5 mg ( labeled ‘a’; Fig. 4A). Targeted DR % was calculated with respect to 56 mg offered in the ‘no DR’ group. Based on actual food consumed (labeled ‘b’; Fig. 4A), the achieved DR % was calculated with respect to 43 mg, the mean food consumed by flies in the ‘no DR’ group (Fig. 4A).

#### vii) TRF feeding protocol with DIETS

Groups of 20 male flies each were split into three groups. Two *ad libitum* feeding groups, where flies were switched to fresh food vials either every 24 h [ALF (24h)] or every 12 h [ALF (12+12)] were set. A third time-restricted feeding (TRF) group was offered food only during the ‘lights on’ phase (ZT = 0 - 12). TRF flies were switched on DIETS vials with no cups during the ‘lights off’ phase (ZT = 12 - 24). All groups were habituated similarly to their experimental conditions (schema; Fig. 4B). Food consumption was measured whenever flies were switched to fresh vials as indicated (schema, Fig. 4B).

### E. Starvation resistance assay

To compare the starvation resistance of flies, we allowed flies in varying group sizes (10, 25 and 50 males) to feed from DIETS vial and regular food vial for 1 day and 7 days, following two days of usual habituation. For 7 days of feeding in DIETS vials, flies were transferred to new DIETS vials every 24 h, while normal vials were changed on alternative days. Flies were then transferred to 0.75 - 1 % agar starvation vials to measure starvation resistance. Dead flies were counted every 24 h. The survival time of individual flies was recorded after week-long feeding on HSD or sHFD. The starvation resistance of flies was computed by the Kaplan Meier plot using graphpad prism.

### F. Gut lipid Oil-Red-O staining

For midgut-deposited fat droplet staining, the fly’s abdomen was dissected in 1X PBS with fine forceps (55-INOX, FST) to remove the gut. Gut tissues were fixed in 4 % paraformaldehyde (prepared from pre-dissolved solution) (Sigma-Aldrich: P6148-500G) for 20 minutes at room temperature on a rotary shaker. Dissected guts are then rinsed twice for 10 minutes each with 1X PBS to remove the fixative. Fixed guts were then incubated in Oil-Red-O stain for 30 minutes. Oil-Red-O stains were prepared by dissolving 10 mg Oil-Red-O (powder) in 10 mL isopropanol (0.1 % Oil-Red-O). A 6:4 ratio of 0.1 % Oil-Red-O in isopropanol to 4 mL distilled water was prepared and passed through a 0.22 □m syringe filter (Merck: Z260479) before use. The stained gut tissues were rinsed twice in distilled water for 10 minutes each. Stained guts were then mounted on a glass slide, and images were captured using an upright Nikon fluorescent microscope (Nikon Eclipse 80i) with a 20X objective (Plan Fluor 20X, 0.50 NA Ph1 DLL ∞/0.17 WD 2.1).

### G. RNA extraction and qPCR

After habituation on CD for 2 days, male flies were transferred on either 20 % HFD or CD for 1, 4, and 7 days in DIETS vials and then flash-frozen and stored at −80 C. For RNA extraction, 30 files were vortex-decapitated, and heads were collected on dry ice, lysed in Trizol (Invitrogen: 15596026), and total RNA was extracted using a Zymo Direct Zol kit (Cat No: R2051), and RNA was purified using a Zymo RNA Clean and Concentrator kit (Cat No: R1013) as per the manufacturer’s instructions. After RNA extraction, cDNA was prepared using a High-Capacity cDNA Reverse Transcription kit (Thermo-Fisher Scientific, 4368814). qPCR was performed using KAPA SYBR FAST Universal (Sigma Aldrich, KK4602) on a QuantStudio™ 6 Flex Real-Time PCR System (Applied Biosystems, 4485691). Fold change for each gene was determined using the 2^UVV:.^ method (Schmittgen and Livak 2008).

To obtain a comprehensive list of DEGs upon exposure to sHFD, we re-analysed the raw data generated by a previous study (Stobdan et al. 2019) using DESeq2 (Love, Huber, and Anders 2014) and used p-adjusted value of <0.05 as cut-off. Full analysis results are included in Table ST3. From this analysis, we selected top three of the most upregulated genes-*takeout*, *Obp83a*, and *Cyp4e3*, and one downregulated gene-*TotA*. The primers used were as follows: For RPL32 (endogenous control: FBgn0002626): RPL32-F: CGCCACCAGTCGGATCGATAT and RPL32-R: CATGTGGCGGGTGCGCTTGTT. For TotA (FBgn0028396), TotA-F: CTCTTATGTGCTTTGCACTGCTG and TotA-R: TTTTGGAGTCATCGTCCTGGG. For Cyp4e3 (FBgn0015035), Cyp4e3-F: CCCGATGTCCAGAGGAAATTGT and Cyp4e3-R: GCGCTGCGCCTCCTTAATA. For takeout (FBgn0039298), to-F: CAGCGGAACTGGACAGAGTAAC and to-R: GCGCCTTGTCCCCATTGAACA. For Obp83a (FBgn0011281), Obp83a-F: CGAGGACGAGAAGCTCAAATGC and Obp83a-R: CGGATGGACGCAGCCCTTGG.

### H. DIETS-arena and video analysis

We developed a flat arena setup, DIETS-arena, with an easily replaceable 3D printed food cup (transparent PLA material) in the center to enable temporal and spatial analysis of flies along with food consumption measurement. Each arena measured 6.5* 6.5 * 0.4 cm, and the whole setup comprised of six arenas (Fig. S6A; left panel). This design was adapted from a previous study (Charroux and Royet 2020). 0 - 4 day old male flies were sorted under CO_2_ anesthesia in groups of 15 - 20 and habituated in DIETS vial with CD for 2 days. Arenas were cleaned with warm water and wiped clean thoroughly with a microfibre cloth. 500 □L of 1 % agar was pipetted down in a ring form at a distance of 1 cm away from the edge of the central food cup (Fig. S6A; left panel). The agar was left to dry for 1 h, and subsequently, the arenas were covered with a transparent acrylic sheet. The initial weight of the food cup with CD was weighed and was immediately placed into the slot provided at the center of the arena to prevent evaporation. To transfer flies, the whole setup was inverted. Food cups from individual arenas were taken out one at a time, flies were transferred from a vial to the arena via a custom-designed funnel, and food cups were placed back immediately to prevent escape. The arena was then placed in the image capture setup (Fig. S6A; right panel). Arenas without flies were maintained alongside as evaporation controls. This experiment was initiated at ZT = 1. Images were captured via Arducam HQ IR-CUT camera with a 6 mm CS lens (Cat. 1251599, from robu.in), connected to a Raspberry Pi 4B. The Libcamera package was used to capture time-lapse images at 0.2 fps. The captured images were then undistorted using the Computer Vision Toolbox in MATLAB R2022b. These images were later compressed into videos of 10 fps using ImageJ ( version 1.53t). A previously published MATLAB code (Charroux and Royet 2020) was modified to quantify the number of flies in a defined area.

To determine the number of flies on or near the food cup, we defined a circular zone around the cup (Food_ROI_), at a distance of 90 pixels from the center of the food cup. This figure was a summation of the longest male body length (40 pixels: data not shown) and the food cup radius (50 pixels; see Fig. S6A). As it sometimes proved difficult to tell flies apart in the Food_ROI_, especially when many were present, we applied an inverted logic to determine the number of flies in each captured frame outside the Food_ROI_ in each arena. This ultimately allowed us to calculate the % occupancy of Food_ROI_ per frame as :

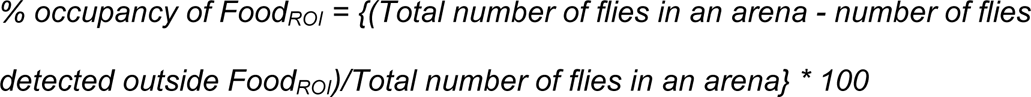

### I. Data plotting and statistics

GraphPad Prism version 8.3.1 was used for most main data plotting. Almost all data are represented as scatter plots with error bars representing the mean with a 95 % confidence interval (CI). Starvation resistance was plotted as Kaplan-Meier graph. Estimation statistics, including effect sizes, the margin of error (MOE), and relevant comparisons between groups performed for each experiment are reported in Supplementary Table 2 (ST2). For all figures, unless mentioned, the effect size is calculated as the mean difference between the two groups. This represents the magnitude of difference between the two groups. The effect size is represented as the dot in the spread. The two black lines that extend from the effect size dot denote the 95% CI and are a measure of the precision of the data. To generate the CI, around 5000 bias-corrected and accelerated (BCa) bootstrap resampling was done and the effect size of each resample was plotted to generate the CI arms. The magnitude of one arm of the confidence interval width represents the MOE. The effect size for Fig. 1E and 2A was measured by the standardized mean difference, Hedge’s g. Hedge’s g is the difference in mean of two groups, standardized over the pooled standard deviation of the two groups, with a correction applied for small (∼ less than 20) group sizes (Turner and Bernard 2006). All estimation/delta plots were generated using an online website: www.estimationstats.com (Ho et al. 2019). We extracted data for EX-Q and CAFE from (Wu et al. 2020), Fig. 3C, using WebPlotDigitizer (<u>automeris.io/WebPlotDigitizer</u>).

## Acknowledgment

We thank all members of the brain and feeding behavior laboratory, NCCS, for their valuable insights in the synthesis of the manuscript. We would like to thank Radhika Mohandasan for early reviewing and aiding in the submission of the manuscript and Asmita Dogra for estimating the weight of fly eggs and Mohd. Tanzeel Aalam for helping with the video. We thank Deevitha Balasubramanian for testing the earliest versions of the DIETS assay. We thank the rest of our labs for their feedback on the manuscript. We thank our lab technician Rajkumar Pawar for managing lab purchases and preparing fly food. We thank Umesh Chachala and Jagtap for lab maintenance. MRT was funded by a Department of Biotechnology (DBT) doctoral scholarship. PC was funded by the University Grants Commission (UGC) and supported by a Ramanujan Fellowship award to GD (SB/S2/RJN-048/2017). SS was hired as a project JRF from a Core Research Grant (CRG/2019/005587) from SERB, DST. GoI. BP is funded by the Council of Scientific and Industrial Research (CSIR) doctoral scholarship. RSPY is supported by a TMA Pai fellowship from MAHE. PA lab is funded by the Ramalingaswami Re-entry Fellowship (BT/RLF, Re-entry/34/2018), and DBT, Research grant (BT/PR36166/BRB/10/1859/2020) by the Department of Biotechnology (DBT), Ministry of Science and Technology, Government of India. GD was a Ramanujan Fellow (SB/S2/RJN-048/2017) awarded by SERB. The GD lab is also funded by a Core Research Grant (CRG/2019/005587) from SERB and generous intramural funding support from NCCS.

## Contributions

G.D. and M.R.T. and P.C. conceived the design of the DIETS assay. G.D. conceived the DIETS-arenas assay. S.S. built the DIETS-arenas and modified the code for DIETS-arena from work cited. Initial trials and design refinement was done by G.D, M.R.T, P.C, S.S, and B.P. Experiments on starvation resistance following HSD or sHFD were carried out by B.P. qPCR experiments were done by R.S.P.Y performed, and supervised by P.A. Both R.S.P.Y and P.A analyzed the data. All experiments, data analysis, and plotting in the manuscript were performed by G.D, P.A, M.R.T, P.C, B.P., S.S, and R.S.P.Y. Statistical analysis was performed by B.P and S.S. G.D directed the research and wrote the manuscript with inputs from all.

