## Supplement File 1. S1 - S6, ST 1-2 for "DIETS: a simple and sensitive assay to measure and control the intake of complex solid foods, like high-fat diets, in *Drosophila*"

### Supplementary Figure 1.

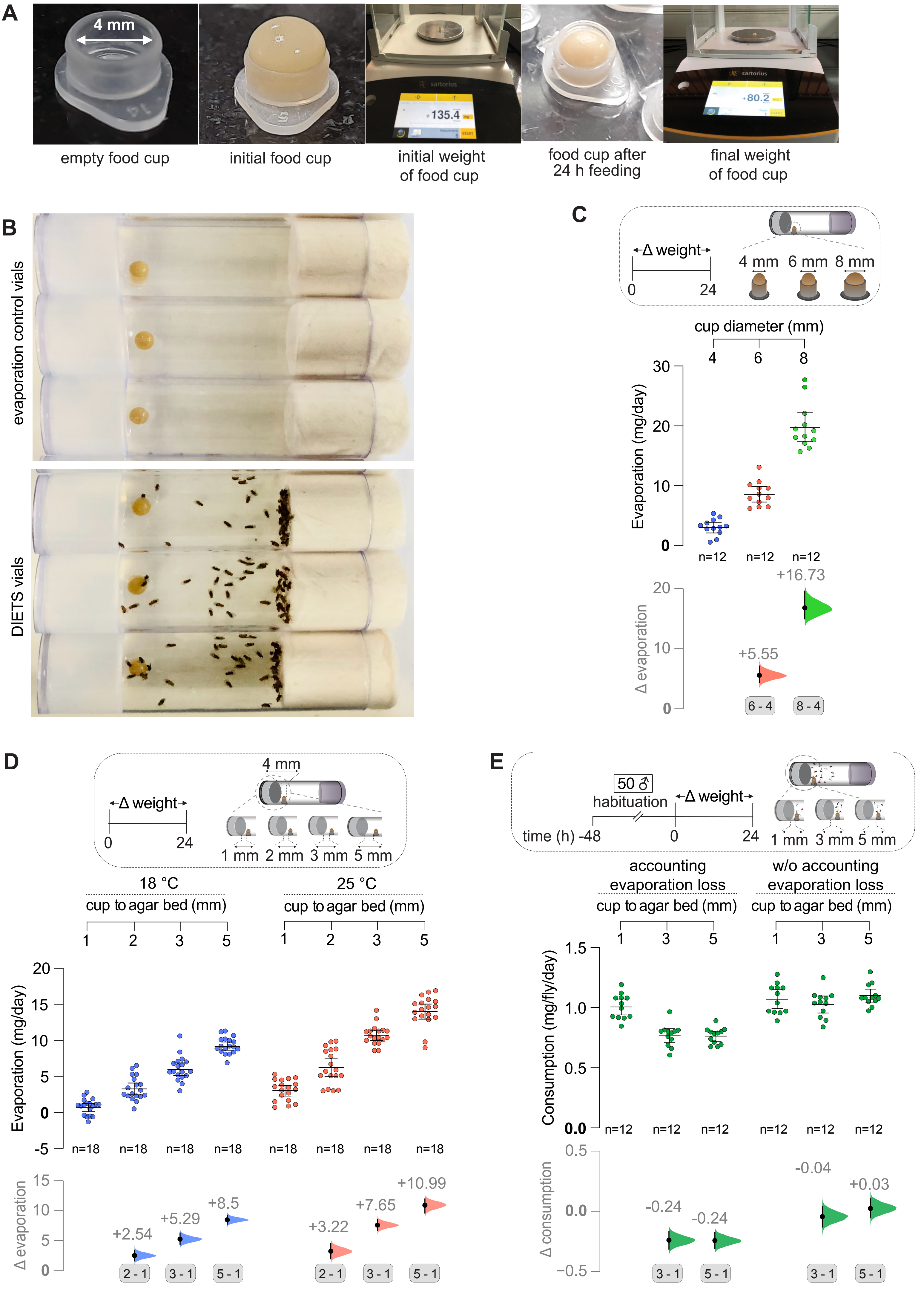

**Supplementary Figure 1. Minimizing food weight loss by evaporation in DIETS (with Figure 1).**

**(A)** Representative images of food cups before and after a DIETS experiment with 50 male flies. Also, an image of the analytical weighing balance showing typical range of weight change when 50 male flies have fed for 24 h. **(B)** Representative image of evaporation vials without flies and DIETS vials. **(C)** Evaporation loss from food cups of different diameters was measured over 24 h without flies. Evaporation loss from food cup increases with increasing food cup diameter. **(D)** Evaporation loss of CD weight was measured over 24 h at 18 °C and 25 °C, over varying food cup to agar bed distance. Evaporation loss increased with an increase in temperature and food cup to agar bed distance. **(E)** The consumption of CD by 50 male flies was measured over 24 h at 25 °C, while varying the distance between the food cup and the agar bed. Evaporation control vials without flies were prone to underestimate feeding, especially when cups were placed further away from the agar bed. Scatter plots of raw data with mean ± 95 % CI are shown. Δ 'effect size' plots below each raw data graph depict mean differences (black dots; values labeled), between the two relevant groups being compared. The 95 % CI (black lines) and distribution of the mean differences (curve), generated by bootstrap resampling of the data are also shown. See Materials and Methods for further details.

### Supplementary Figure 2.

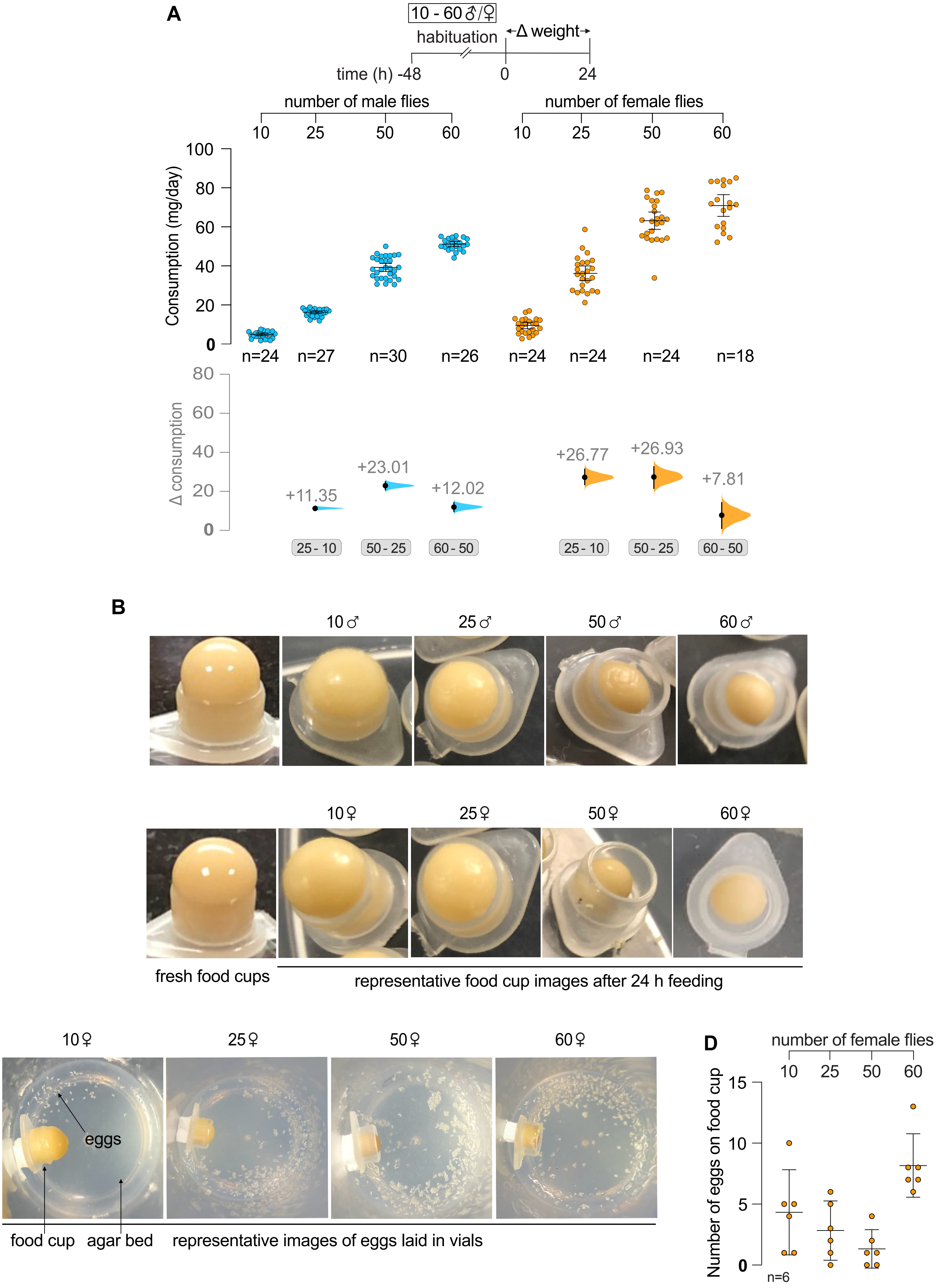

**Supplementary Figure 2. Feeding and egg laying in DIETS vials (with Figure 1).** **(A)** Total consumption per group is plotted for Figure 1B. As the group size increases, the total food consumed also increases. **(B)** Representative food cups before and after 24 h consumption by different group sizes. **(C)** Eggs laid on the agar bed in a 24 h period inside DIETS vials **(D)** Quantification of the number of eggs laid on the food cup in DIETS vials in the same period. Scatter plots of raw data with mean ± 95 % CI are shown. Δ 'effect size' plots below each raw data graph depict mean differences (black dots; values labeled), between the two relevant groups being compared. The 95 % CI (black lines) and distribution of the mean differences (curve), generated by bootstrap resampling of the data are also shown. See Materials and Methods for further details.

**Supplementary Figure 3.**

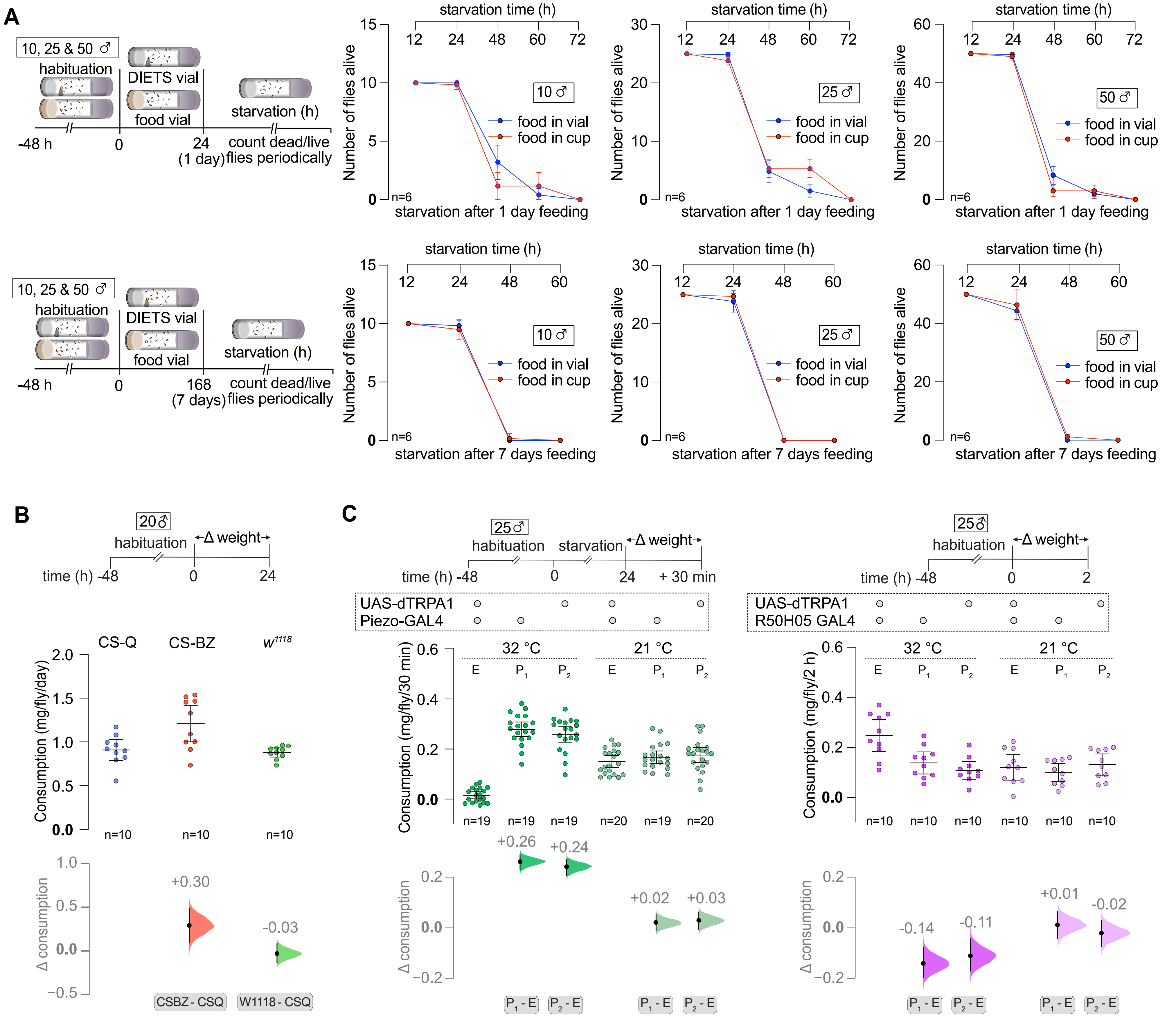

**Supplementary Figure 3. Flies are not starving after being in DIETS vials. DIETS is compatible with thermogenetic manipulations (with Figure 1).**

**(A)** Starvation resistance of flies in different group sizes measured after 1 d and 7 d feeding in DIETS and regular food vials. No evident change in starvation resistance was observed. (B) 24 h CD consumption rate was measured with three different fly strains in the lab. (C) 30 minutes CD consumption was measured upon TRPA1-induced activation of Piezo neurons in 24 h starved flies. Piezo-GAL4 driven activation of neurons with UAS-dTRPA1 inhibits feeding in starved flies compared to parental controls and temperature controls (left panel). 2 h CD consumption was measured upon TRPA1-induced activation of R50H05 neurons in sated flies. Artificial activation of R50H05-GAL4 neurons increases feeding in sated flies compared to parental controls and temperature controls (right panel). These results show the compatibility of DIETS with thermogenetic manipulations at 32 °C. Parental controls and respective temperature controls at 21 °C are shown for both. Scatter plots of raw data with mean ± 95 % CI are shown. E represents experimental cross while P1 and P2 represent parental controls. Δ 'effect size' plots below each raw data graph depict mean differences (black dots; values labeled), between the two relevant groups being compared. The 95 % CI (black lines) and distribution of the mean differences (curve) generated by bootstrap resampling of the data are also shown. See Materials and Methods for further details.

### Supplementary Figure 4.

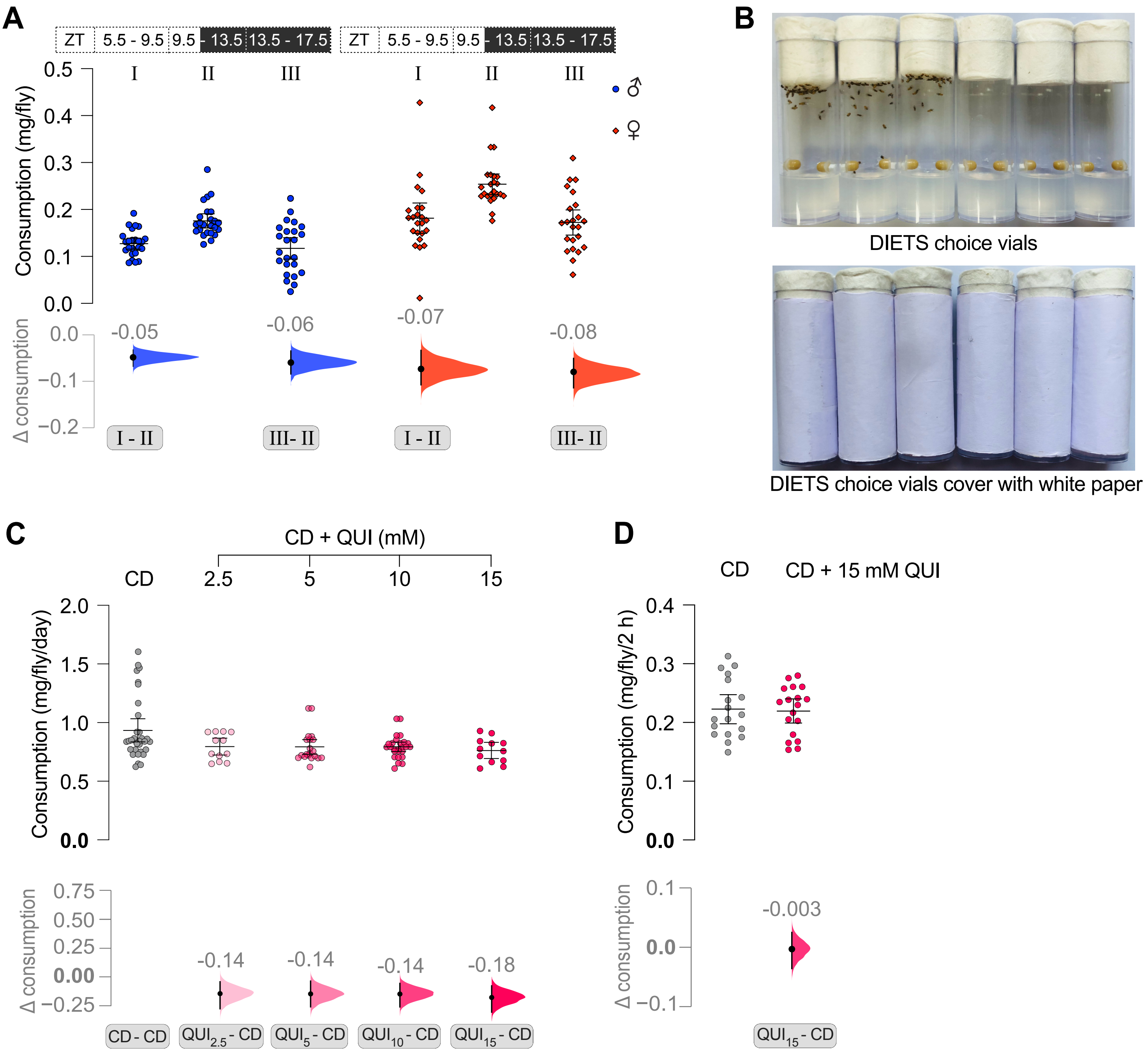

**Supplementary Figure 4. Highlighting evening feeding peak and total consumption in choice setup (with Figure 2).**

**(A)** Cumulative feeding measurements from 2 data points around lights-off time (ZT = 9.5 - 13.5) compared with combined measurements on both sides; from ZT = 5.5 - 9.5 and ZT = 13.5 - 17.5. Based on the data from Fig. 2B. Highlights increased feeding or the presence of a feeding peak around lights-off time for both males and females. **(B)** DIETS choice vials include two 4 mm diameter food cups placed diametrically opposite each other, equidistant from the agar bed. Vials were covered with white paper to provide diffused light and prevent light bias. Choice vials were kept horizontally in incubators. **(C-D)** Combined consumption from both the cups, from respective choice groups used in Figure 3D and 3E, respectively. The total food intake in different choice groups remains largely unaltered. Scatter plots of raw data with mean ± 95 % CI are shown. Δ 'effect size' plots below each raw data graph depict mean differences (black dots; values labeled) between the two relevant groups being compared. The 95 % CI (black lines) and distribution of the mean differences (curve) generated by bootstrap resampling of the data are also shown. See Materials and Methods for further details.

### Supplementary Figure 6.

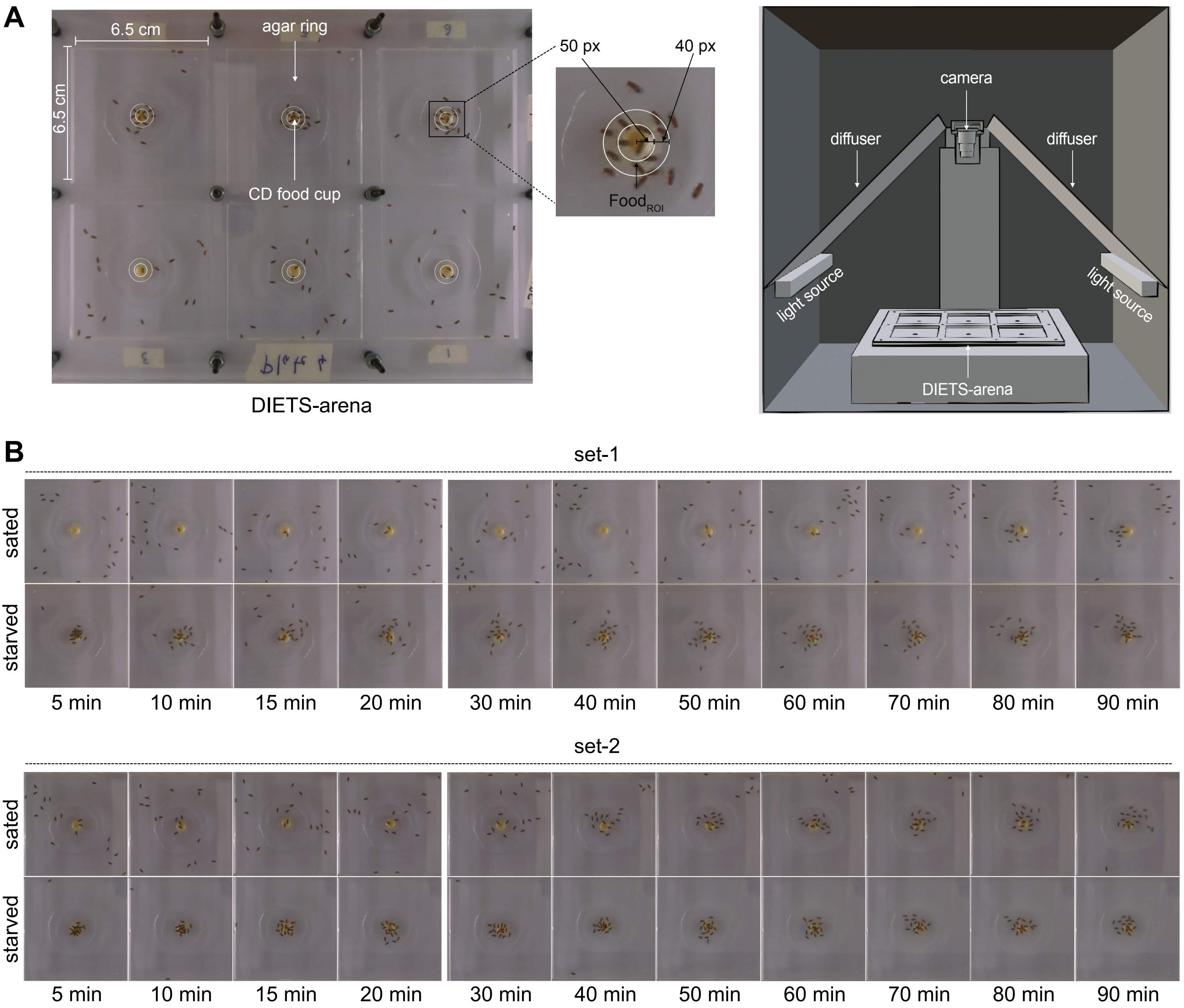

**Supplementary Figure 5. DIETS-arena setup (with Figure 3).**

**(A)** Image showing DIETS-arenas and associated imaging setup. Pertinent features are marked. Inset highlights a $Food_{ROI}$ marked around a food cup. **(B)** Timeline showing images of representative DIET-arenas at different time points from two different sets of starved and sated flies.

### Supplementary Figure 6.

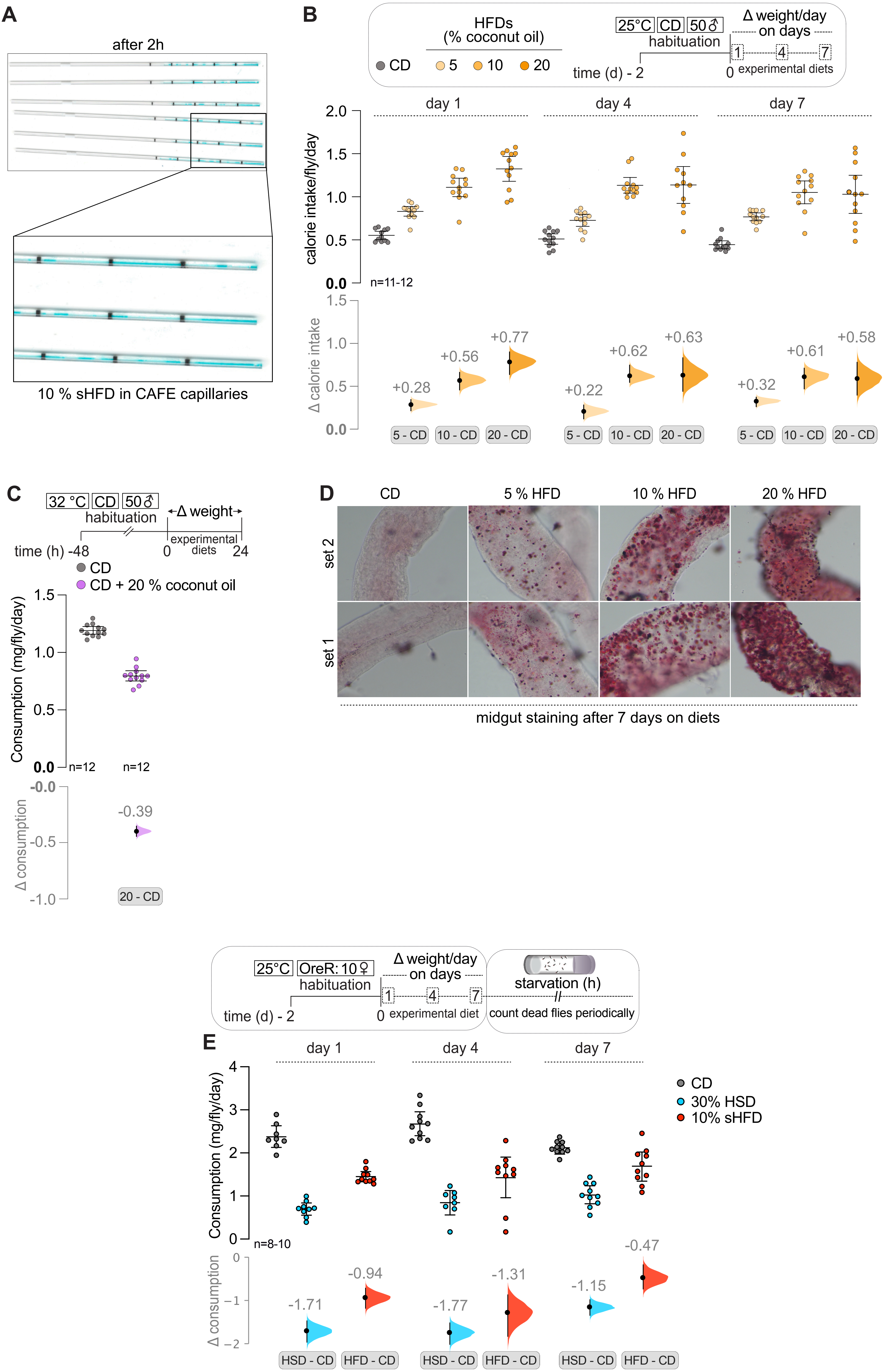

**Supplementary Figure 6. Calorie intake and the suitability of DIETS for high-fat diet intake measurements at higher temperatures (with Figure 3 and 4).**

**(A)** Scan images of capillaries used for CAFE assay with liquid 10% HFD. We noted an intermittent flow of liquid food within the capillaries, which may pose a challenge for utilizing complex diets in the CAFE assay. **(B)** The corresponding calorie intake on HFDs from Figure 3A is depicted. Calorie intake seems to plateau at 10 % coconut oil in the food. **(C)** Feeding was measured on a control diet (CD) and a 20 % saturated HFD at 32 °C. Such measurements would likely be virtually impossible any other way than offering food in small cups. Shows that DIETS is suitable for measuring feeding on fatty food even at higher temperatures. **(D)** Oil-Red-O staining of lipid droplets was performed on the midgut of flies fed different HFDs for 7 days. Fat deposition increased with % of coconut oil in the food. **(E)** 24 h food intake of OreR flies were determined on days 1, 4, and 7 on respective diets. Compared to the control group, flies on sHFD and HSD exhibit reduced feeding. Scatter plots of raw data with mean ± 95 % CI are shown. Δ ‘effect size’ plots below each raw data graph depict mean differences (black dots; values labeled), between the two relevant groups being compared. The 95 % CI (black lines) and distribution of the mean differences (curve) generated by bootstrap resampling of the data are also shown. See Materials and Methods for further details.

| Supplementary Table 1 | | | |
|---|---|---|---|
| Ingredients for 100 mL diet preperation | | | |
| | control diet (CD) | 30 % HSD | 10 % sHFD |
| Agar | 1 g | 1 g | 1 g |
| Brewer's yeast | 8 g | 8 g | 8 g |
| Yeast extract | 2 g | 2 g | 2 g |
| Peptone | 2 g | 2 g | 2 g |
| Sucrose | 5.1 g | 30 g | 5.1 g |
| Coconut oil | - | - | 10 g |
| $MgSO_4$ X $6H_2O$ | 200 ul | 200 ul | 200 ul |
| $CaCl_2$ X $2H_2O$ | 300 ul | 300 ul | 300 ul |
| Propionic acid | 600 ul | 600 ul | 600 ul |
| Methyl-4-hydrobenzoate (1.19 g in 54 mL 95 % ethanol) | 1000 ul | 1000 ul | 1000 ul |

| Supplementary Table 2 | | | | | | | | |
|---|---|---|---|---|---|---|---|---|
| **Figure** | **Unit** | **Group** | **Mean** | **SD** | **n** | **Relevant comparison of groups** | **Mean difference ES** | **MOE (1/2 CI)** |
| | | 10 ♂ | 0.47 | 0.17 | 24 | ♂ 10 flies vs. ♀ 10 flies | 0.46 | 0.16 |
| | | 25 ♂ | 0.64 | 0.07 | 27 | ♂ 25 flies vs. ♀ 25 flies | 0.8 | 0.14 |
| | | 50 ♂ | 0.82 | 0.12 | 30 | ♂ 50 flies vs. ♀ 50 flies | 0.44 | 0.09 |
| | | 60 ♂ | 0.84 | 0.07 | 26 | ♂ 60 flies vs. ♀ 60 flies | 0.34 | 0.09 |
| | | 10 ♀ | 0.92 | 0.39 | 24 | ♂ 10 flies vs. 25 flies | 0.17 | 0.07 |
| | | 25 ♀ | 1.44 | 0.35 | 24 | ♂ 25 flies vs. 50 flies | 0.18 | 0.05 |
| | | 50 ♀ | 1.26 | 0.21 | 24 | ♂ 50 flies vs. 60 flies | 0.02 | 0.05 |
| Figure 1 B | Consumption (mg/fly/day) | 60 ♀ | 1.18 | 0.19 | 18 | | | |
| | | ♂ in 18 °C | 0.33 | 0.06 | 12 | ♂ in 18 °C vs. 22 °C | 0.23 | 0.05 |
| | | ♂ in 22 °C | 0.56 | 0.06 | 12 | ♂ in 18 °C vs. 25 °C | 0.49 | 0.09 |
| Figure 1 C | Consumption (mg/fly/day) | ♂ in 25 °C | 0.82 | 0.15 | 12 | ♂ in 22 °C vs. 25 °C | 0.26 | 0.09 |
| | | ♂ on 0.5X CD | 1.13 | 0.12 | 12 | ♂ on 1X vs. 0.5X | 0.2 | 0.09 |
| | | ♂ on 0.75X CD | 0.94 | 0.11 | 12 | ♂ on 1X vs. 0.75X | 0.01 | 0.08 |

| | | | | | | | |
|---|---|---|---|---|---|---|---|
| | | ♂ on 1X CD | 0.93 | 0.1 | 12 | ♂ on 1X vs. 1.5X | -0.22 | 0.07 |
| | | ♂ on 1.5X CD | 0.71 | 0.07 | 12 | ♂ on 1X vs. 2X | -0.34 | 0.07 |
| | | ♂ on 2X CD | 0.6 | 0.07 | 12 | ♂ on 0.5X vs. 0.75X | -0.19 | 0.09 |
| Figure 1 D | Consumption (mg/fly/day) | | | | | ♂ on 1.5X vs. 2X | -0.11 | 0.05 |
| | | DIETS: ♀ on 1X YS diet | 1.73 | 0.19 | 10 | DIETS: ♀ on 1X vs. 2X | - 2.59* | 1.09* |
| | | DIETS: ♀ on 2X YS diet | 1.12 | 0.25 | 10 | CAFE: ♀ on 1X vs. 2X | - 3.12* | 1.87* |
| | | DIETS: ♀ on 4X YS diet | 0.46 | 0.18 | 10 | EX-Q: ♀ on 1X vs. 2X | - 5.51* | 1.46* |
| | | CAFE: ♀ on 1X YS diet | 1.65 | 0.27 | 10 | DIETS: ♀ on 2X vs. 4X | - 2.87* | 2.11* |
| | | CAFE: ♀ on 2X YS diet | 0.93 | 0.17 | 10 | CAFE: ♀ on 2X vs. 4X | - 2.39* | 1.52* |
| | | CAFE: ♀ on 4X YS diet | 0.58 | 0.09 | 9 | EX-Q: ♀ on 2X vs. 4X | - 2.98* | 1.78* |
| | | EX-Q: ♀ on 1X YS diet | 2.39 | 0.23 | 10 | | | |
| Figure 1 E | Consumption (mg/fly/day) | EX-Q: ♀ on 2X YS diet | 1.28 | 0.15 | 10 | | | |
| | | EX-Q: ♀ on 4X YS diet | 0.76 | 0.18 | 10 | | | |
| | | ♂ day 1 feeding | 1.01 | 0.15 | 11 | ♂ feeding day 1 vs. day 4 | -0.07 | 0.12 |
| | | ♂ day 4 feeding | 0.94 | 0.15 | 12 | ♂ feeding day 1 vs. day 7 | 0.08 | 0.12 |
| | | ♂ day 7 feeding | 1.1 | 0.16 | 12 | ♂ feeding day 1 vs. day 14 | -0.15 | 0.11 |

| | | | | | | | |
|---|---|---|---|---|---|---|---|
| | | ♂ day 14 feeding | 0.86 | 0.13 | 12 | ♂ feeding day 1 vs. day 21 | -0.04 | 0.14 |
| | | ♂ day 21 feeding | 0.96 | 0.2 | 11 | ♂ feeding day 1 vs. day 28 | -0.27 | 0.24 |
| Figure 1 F | Consumption (mg/fly/day) | ♂ day 28 feeding | 0.74 | 0.38 | 11 | | | |
| | | evaporation on 4 mm cup | 3.13 | 1.41 | 12 | cup diameter of 6 mm vs. 4 mm | 5.55 | 2.775 |
| | | evaporation on 6 mm cup | 8.68 | 2.05 | 12 | cup diameter of 8 mm vs. 4 mm | 16.73 | 8.365 |
| Figure S1C | Evaporation (mg/day) | evaporation on 8 mm cup | 19.87 | 3.8 | 12 | | | |
| | | 18 °C - 1 mm from cup to agar bed | 0.8 | 1.11 | 18 | 18 °C: 1 mm vs. 2 mm distance from agar bed | 2.54 | 0.89 |
| | | 18 °C - 2 mm from cup to agar bed | 3.34 | 1.66 | 18 | 18 °C: 1 mm vs. 3 mm distance from agar bed | 5.29 | 0.94 |
| | | 18 °C - 3 mm from cup to agar bed | 6.09 | 1.72 | 18 | 18 °C: 1 mm vs. 5 mm distance from agar bed | 8.5 | 0.72 |
| | | 18 °C - 5 mm from cup to agar bed | 9.3 | 1.15 | 18 | | | |
| | | 25 °C - 1 mm from cup to agar bed | 3.11 | 1.45 | 18 | 25 °C: 1 mm vs. 2 mm distance from agar bed | 3.22 | 1.27 |

| | | | | | | | | |
|---|---|---|---|---|---|---|---|---|
| | | 25 °C - 2 mm from cup to agar bed | 6.32 | 2.44 | 18 | 25 °C: 1 mm vs. 3 mm distance from agar bed | 7.65 | 0.9 |
| | | 25 °C - 3 mm from cup to agar bed | 10.76 | 1.39 | 18 | 25 °C: 1 mm vs. 5 mm distance from agar bed | 10.99 | 1.18 |
| Figure S1D | Evaporation (mg/day) | 25 °C - 5 mm from cup to agar bed | 14.09 | 2.1 | 18 | | | |
| | | accounting evaporation loss - 1 mm from cup to agar bed | 1.01 | 0.1 | 12 | accounting evaporation loss - 1 mm vs. 3 mm from cup to agar bed | -0.24 | 0.07 |
| | | accounting evaporation loss - 3 mm from cup to agar bed | 0.77 | 0.09 | 12 | accounting evaporation loss - 1 mm vs. 5 mm from cup to agar bed | -0.24 | 0.07 |
| | | accounting evaporation loss - 5 mm from cup to agar bed | 0.77 | 0.07 | 12 | | | |
| | | without accounting evaporation loss - 1 mm from cup to agar bed | 1.07 | 0.12 | 12 | without accounting evaporation loss - 1 mm vs. 3 mm from cup to agar bed | -0.04 | 0.09 |

| | | | | | | | | |
|---|---|---|---|---|---|---|---|---|
| | | without accounting evaporation loss - 3 mm from cup to agar bed | 1.03 | 0.11 | 12 | without accounting evaporation loss - 1 mm vs. 5 mm from cup to agar bed | 0.03 | 0.08 |
| Figure S1E | Evaporation (mg/day) | without accounting evaporation loss - 5 mm from cup to agar bed | 1.1 | 0.09 | 12 | | | |
| | | ♂ - 10 flies | 4.66 | 1.73 | 24 | ♂ 10 flies vs. 25 flies | 11.35 | 0.96 |
| | | ♂ - 25 flies | 16.01 | 1.84 | 27 | ♂ 25 flies vs. 50 flies | 23.01 | 2.09 |
| | | ♂ - 50 flies | 39.02 | 5.55 | 30 | ♂ 50 flies vs. 60 flies | 12.02 | 2.27 |
| | | ♂ - 60 flies | 51.04 | 3.04 | 22 | | | |
| | | ♀ - 10 flies | 9.24 | 3.86 | 24 | ♀ 10 flies vs. 25 flies | 26.77 | 3.85 |
| | | ♀ - 25 flies | 36.01 | 8.85 | 24 | ♀ 25 flies vs. 50 flies | 26.93 | 5.38 |
| | | ♀ - 50 flies | 62.94 | 10.51 | 24 | ♀ 50 flies vs. 60 flies | 7.81 | 6.41 |
| Figure S2A | Consumption (mg/fly/day) | ♀ - 60 flies | 70.75 | 11.15 | 18 | | | |
| | | CS-Q | 0.91 | 0.17 | 10 | food consumption with CS-Q flies vs.. CSBZ flies | 0.3 | 0.19 |

| | | | | | | | | |
|---|---|---|---|---|---|---|---|---|
| Figure S3B | Consumption (mg/fly/day) | CS-Bz | 1.2 | 0.29 | 10 | food consumption with CS-Q flies vs.. w1118 flies | -0.03 | 0.11 |
| | | w1118 | 0.88 | 0.07 | 10 | | | |
| | Consumption (mg/fly/30min) | 32 °C: Piezo-GAL4/UAS - dTRPA1 | 0.02 | 0.03 | 19 | 32 °C: food consumption of E vs. P1 | 0.26 | 0.03 |
| | | 32 °C: Piezo-GAL4 | 0.28 | 0.06 | 19 | 32 °C: food consumption of E vs. P2 | 0.24 | 0.03 |
| | | 32 °C: UAS - dTRPA1 | 0.26 | 0.07 | 19 | | | |
| | | 21 °C: Piezo-GAL4/UAS - dTRPA1 | 0.15 | 0.05 | 20 | 21 °C: food consumption of E vs. P1 | 0.02 | 0.03 |
| | | 21 °C: Piezo-GAL4 | 0.17 | 0.05 | 19 | 21 °C: food consumption of E vs. P2 | 0.03 | 0.04 |
| | | 21 °C: UAS - dTRPA1 | 0.17 | 0.07 | 20 | | | |
| | | 32 °C R50H05-GAL4/UAS - dTRPA1 | 0.25 | 0.09 | 10 | 32 °C: food consumption of E vs. P1 | -0.14 | 0.06 |
| | | 32 °C R50H05-GAL4 | 0.11 | 0.05 | 10 | 32 °C: food consumption of E vs. P2 | -0.11 | 0.06 |
| | | 32 °C UAS - dTRPA1 | 0.14 | 0.06 | 10 | | | |

| | | | | | | | | |
|---|---|---|---|---|---|---|---|---|
| | | 21°C : Piezo-GAL4/UAS - dTRPA1 | 0.12 | 0.07 | 10 | 21 °C: food consumption of E vs. P1 | 0.01 | 0.05 |
| | | 21°C: Piezo-GAL4 | 0.13 | 0.06 | 10 | 21 °C: food consumption of E vs. P2 | -0.02 | 0.05 |
| Figure S3C | Consumption (mg/fly/2h) | 21°C: UAS - dTRPA1 | 0.1 | 0.05 | 10 | | | |
| | | DIETS assay - 0 h starvation | 0.01 | 0.01 | 24 | 0 h vs. 6 h starved | 4.65* | 0.80* |
| | | DIETS assay - 6 h starvation | 0.06 | 0.01 | 18 | 6 h vs. 12 h starved | 1.82* | 0.81* |
| | | DIETS assay - 12 h starvation | 0.1 | 0.02 | 24 | 12 h vs. 24 h starved | 1.35* | 0.69* |
| | Consumption (mg/fly/30min) | DIETS assay - 24 h starvation | 0.13 | 0.03 | 23 | | | |
| | | blue dye assay - 0 h | 0.13 | 0.23 | 24 | 0 h vs. 6 h starved | 1.69* | 0.81* |
| | | blue dye assay - 6 h | 0.48 | 0.16 | 18 | 6 h vs. 12 h starved | 0.87* | 0.58* |
| | | blue dye assay - 12 h | 0.63 | 0.17 | 24 | 12 h vs. 24 h starved | 0.75* | 0.64* |
| Figure 2A | Absorbance at 630nm | blue dye assay - 24 h | 0.81 | 0.29 | 23 | | | |
| | | cup A1: CD | 0.46 | 0.16 | 32 | cup A1: CD vs. cup B1: CD | 0.01 | 0.07 |
| | | cup A2: CD | 0.39 | 0.04 | 12 | cup A2: CD vs. cup B2: CD + QUI ( 2.5 mm) | 0.01 | 0.06 |

| | | | | | | | | |
|---|---|---|---|---|---|---|---|---|
| | | cup A3: CD | 0.42 | 0.08 | 20 | cup A3: CD vs. cup B3: CD + QUI ( 5 mm) | -0.05 | 0.05 |
| | | cup A4: CD | 0.47 | 0.1 | 26 | cup A4: CD vs. cup B4: CD + QUI (10 mm) | -0.14 | 0.04 |
| | | cup A5: CD | 0.51 | 0.1 | 12 | cup A5: CD vs. cup B5: CD + QUI (15 mm) | -0.26 | 0.07 |
| | | cup B1: CD | 0.48 | 0.14 | 32 | | | |
| | | cup B2: CD + QUI (2.5 mm) | 0.4 | 0.11 | 12 | | | |
| | | cup B3: CD + QUI (5 mm) | 0.37 | 0.08 | 20 | | | |
| | | cup B4: CD + QUI (10 mm) | 0.32 | 0.07 | 26 | | | |
| Figure 2C | Consumption (mg/fly/day) | cup B5: CD + QUI (15 mm) | 0.25 | 0.09 | 12 | | | |
| | | cup A1: CD | 0.12 | 0.03 | 18 | cup A1: CD vs. cup B1: CD | -0.01 | 0.02 |
| | | cup A2: CD | 0.11 | 0.03 | 18 | cup A1: CD vs. cup B2: CD + QUI (15 mm) | -0.09 | 0.02 |
| | | cup B1: CD | 0.16 | 0.03 | 18 | | | |
| Figure 2D | Consumption (mg/fly/ 2 h) | cup B2: CD + QUI (15 mm) | 0.06 | 0.03 | 18 | | | |

| | | | | | | | | |
|---|---|---|---|---|---|---|---|---|
| | | ♂ (I) | 0.13 | 0.03 | 24 | ♂ (II) vs. ♂(I) | -0.05 | 0.02 |
| | | ♂ (II) | 0.18 | 0.04 | 24 | ♂ (II) vs. ♂(III) | -0.06 | 0.03 |
| | | ♂ (III) | 0.12 | 0.05 | 24 | | | |
| | | ♀ (I) | 0.18 | 0.07 | 23 | ♀ (II) vs. ♀(I) | -0.07 | 0.04 |
| | | ♀ (II) | 0.25 | 0.05 | 23 | ♀ (II) vs. ♀(III) | -0.08 | 0.03 |
| Figure S4A | Consumption (mg/fly) | ♀ (III) | 0.17 | 0.06 | 23 | | | |
| | | CD vs. CD | 0.94 | 0.27 | 32 | total consumption in CD - CD vs. CD - QUI (2.5) | -0.14 | 0.12 |
| | | CD vs. CD + QUI (2.5) | 0.8 | 0.11 | 12 | total consumption in CD - CD vs. CD - QUI (5.0) | -0.14 | 0.11 |
| | | CD vs. CD + QUI (5) | 0.79 | 0.13 | 20 | total consumption in CD - CD vs. CD - QUI (10.0) | -0.14 | 0.1 |
| | | CD vs. CD + QUI (10) | 0.79 | 0.1 | 26 | total consumption in CD - CD vs. CD - QUI (15.0) | -0.18 | 0.1 |
| Figure S4C | Consumption (mg/fly/day) | CD vs. CD + QUI (15) | 0.76 | 0.11 | 12 | | | |
| | | CD | 0.22 | 0.05 | 18 | 2 h consumption in CD vs. CD +15 mm QUI | -0.09 | 0.02 |
| Figure S4D | Consumption (mg/fly/2h) | CD + QUI (15) | 0.22 | 0.04 | 18 | | | |

| | | | | | | | | |
|---|---|---|---|---|---|---|---|---|
| | | No DR | 42.76 | 1.07 | 12 | ♂ on 10 % DR vs. no DR (II-I) | -4.58 | 0.67 |
| | | 10 % DR | 38.18 | 0.57 | 12 | ♂ on 20 % DR vs. no DR (III-I) | -6.61 | 0.78 |
| | | 20 % DR | 36.15 | 0.94 | 12 | ♂ on 30 % DR vs. no DR (IV-I) | -11.08 | 0.92 |
| | | 30 % DR | 31.69 | 1.33 | 12 | ♂ on 40 % DR vs. no DR (V-I) | -14.24 | 0.72 |
| | | 40 % DR | 28.52 | 0.79 | 12 | ♂ on 50 % DR vs. no DR (VI-I) | -17.16 | 0.74 |
| Figure 3A | Consumption (mg/day) | 50 % DR | 25.6 | 0.86 | 12 | | | |
| | | TRF | 0.66 | 0.14 | 12 | consumption on ALF (12+12) vs. TRF | 0.06 | 0.14 |
| | | ALF (24) | 0.72 | 0.16 | 12 | consumption on ALF vs. TRF | 0.06 | 0.12 |
| | | ALF (12+12) | 0.72 | 0.21 | 12 | consumption on (a) vs. (b) | 0.06 | 0.08 |
| | | ALF (a) | 0.33 | 0.1 | 12 | | | |
| Figure 3B | Consumption (mg/fly/day) | ALF (b) | 0.38 | 0.16 | 12 | | | |
| | | DIETS vial - sated | 0.07 | 0.03 | 10 | DIETS: sated vs. starved | 0.21 | 0.106191 |
| | | DIETS vial - starved | 0.29 | 0.04 | 11 | DIETS-arena : sated vs. starved | 0.22 | 0.111683 |

| | | | | | | | | |
|---|---|---|---|---|---|---|---|---|
| | | DIETS-arena - sated | 0.21 | 0.05 | 9 | | | |
| Figure 3C | Consumption (mg/fly/2h) | DIETS-arena - starved | 0.43 | 0.05 | 9 | | | |
| | | Day 1 - CD | 0.9 | 0.12 | 11 | Day 1: consumption in CD vs. 5 % HFD | -0.09 | 0.08 |
| | | Day 1 - CD + 5 % oil | 0.81 | 0.09 | 12 | Day 1: consumption in CD vs. 10 % HFD | -0.13 | 0.09 |
| | | Day 1 - CD + 10 % oil | 0.77 | 0.12 | 12 | Day 1: consumption in CD vs. 20 % HFD | -0.32 | 0.09 |
| | | Day 1 - CD + 20 % oil | 0.58 | 0.1 | 12 | | | |
| | | | | | | Day 4: consumption in CD vs. 5 % HFD | -0.13 | 0.1 |
| | | Day 4 - CD | 0.83 | 0.15 | 12 | Day 4: consumption in CD vs. 10 % HFD | -0.05 | 0.1 |
| | | Day 4 - CD + 5 % oil | 0.7 | 0.11 | 12 | Day 4: consumption in CD vs. 20 % HFD | -0.33 | 0.12 |
| | | Day 4 - CD + 10 % oil | 0.78 | 0.1 | 12 | | | |
| | | Day 4 - CD + 20 % oil | 0.5 | 0.14 | 11 | Day 7: consumption in CD vs. 5 % HFD | 0.02 | 0.07 |

| | | | | | | | | |
|---|---|---|---|---|---|---|---|---|
| | | | | | | Day 7: consumption in CD vs. 10 % HFD | 0 | 0.1 |
| | | Day 7 - CD | 0.73 | 0.11 | 12 | Day 7: consumption in CD vs. 20 % HFD | -0.28 | 0.1 |
| | | Day 7 - CD + 5 % oil | 0.75 | 0.07 | 12 | | | |
| | | Day 7 - CD + 10 % oil | 0.73 | 0.14 | 12 | | | |
| Figure 4A | Consumption (mg/fly/day) | Day 7 - CD + 20 % oil | 0.45 | 0.15 | 12 | | | |
| | | Day 1 - CD | 1.46 | 0.18 | 8 | Day 1: energy intake in CD vs. 30% HSD | -0.52 | 0.19 |
| | | Day 1 - 30% HSD | 0.94 | 0.24 | 9 | Day 1: consumption in CD vs. 10% sHFD | 0.64 | 0.18 |
| | | Day 1 - 10% sHFD | 2.11 | 1.64 | 10 | | | |
| | | Day 4 - CD | 1.64 | 0.22 | 10 | Day 4: energy intake in CD vs. 30% HSD | -0.5 | 0.32 |
| | | Day 4 - 30% HSD | 1.14 | 0.44 | 8 | Day 4: energy intake in CD vs. 10% sHFD | 0.43 | 0.56 |
| | | Day 4 - 10% sHFD | 2.07 | 0.91 | 10 | | | |

| | | | | | | | | |
|---|---|---|---|---|---|---|---|---|
| | | Day 7 - CD | 1.3 | 0.09 | 10 | Day 7: energy intake in CD vs. 30% HSD | 0.07 | 0.23 |
| | | Day 7 - 30% HSD | 1.38 | 0.37 | 10 | Day 7: energy intake in CD vs. 10% sHFD | 1.15 | 0.35 |
| Figure 4C | Energy intake fly/day | Day 7 - 10% sHFD | 2.46 | 0.6 | 10 | | | |
| | | CD | 36.26 | 6.61 | 117 | CD vs 30% HSD | 44.39 | 4.09 |
| | | 30% HSD | 80.65 | 17.4 | 111 | CD vs 10% sHFD | 6.26 | 3.66 |
| Figure 4D | Survival time (hours) | 10% sHFD | 42.51 | 8.24 | 105 | | | |
| | | Day 1 - CD | 0.55 | 0.07 | 11 | Day 1: energy intake in CD vs. 5 % HFD | 0.28 | 0.06 |
| | | Day 1 - CD + 5 % oil | 0.83 | 0.09 | 12 | Day 1: energy intake in CD vs. 10 % HFD | 0.56 | 0.1 |
| | | Day 1 - CD + 10 % oil | 1.11 | 0.17 | 12 | Day 1: energy intake in CD vs. 20 % HFD | 0.77 | 0.13 |
| | | Day 1 - CD + 20 % oil | 1.33 | 0.23 | 12 | | | |
| | | | | | | Day 4: energy intake in CD vs. 5 % HFD | 0.22 | 0.08 |
| | | Day 4 - CD | 0.51 | 0.09 | 12 | Day 4: energy intake in CD vs. 10 % HFD | 0.62 | 0.1 |
| | | Day 4 - CD + 5 % oil | 0.73 | 0.11 | 12 | Day 4: energy intake in CD vs. 20 % HFD | 0.63 | 0.19 |

| | | | | | | | |
|---|---|---|---|---|---|---|---|
| | | Day 4 - CD + 10 % oil | 1.13 | 0.14 | 12 | | |
| | | Day 4 - CD + 20 % oil | 1.14 | 0.32 | 11 | Day 7: energy intake in CD vs. 5 % HFD | 0.32 | 0.05 |
| | | | | | | Day 7: energy intake in CD vs. 10 % HFD | 0.61 | 0.12 |
| | | Day 7 - CD | 0.45 | 0.07 | 12 | Day 7: energy intake in CD vs. 20 % HFD | 0.58 | 0.19 |
| | | Day 7 - CD + 5 % oil | 0.77 | 0.07 | 12 | | | |
| | | Day 7 - CD + 10 % oil | 1.05 | 0.21 | 12 | | | |
| Figure S6A | Energy intake/fly/day | Day 7 - CD + 20 % oil | 1.03 | 0.35 | 12 | | | |
| | | 32 °C Day 1 - CD | 1.19 | 0.05 | 12 | 32 °C: consumption of CD vs. 20 % HFD | -0.39 | 0.05 |
| Figure S6B | Consumption (mg/fly/day) | 32 °C Day 1 - CD + 20 % oil | 0.79 | 0.07 | 12 | | | |

*** Effect size calculated as standardized mean difference: Hedge's g**
